# The evolution of widespread recombination suppression on the Dwarf Hamster (*Phodopus*) X chromosome

**DOI:** 10.1101/2021.11.15.468705

**Authors:** Emily C. Moore, Gregg W. C. Thomas, Sebastian Mortimer, Emily E. K. Kopania, Kelsie E. Hunnicutt, Zachary J. Clare-Salzler, Erica L. Larson, Jeffrey M. Good

## Abstract

The mammalian X chromosome shows strong conservation among distantly related species, limiting insights into the distinct selective processes that have shaped sex chromosome evolution. We constructed a chromosome-scale *de novo* genome assembly for the Siberian dwarf hamster (*Phodopus sungorus*), a species reported to show extensive recombination suppression across an entire arm of the X chromosome. Combining a physical genome assembly based on shotgun and long-range proximity ligation sequencing with a dense genetic map, we detected widespread suppression of female recombination across ∼65% of the *Phodopus* X chromosome. This region of suppressed recombination likely corresponds to the Xp arm, which has previously been shown to be highly heterochromatic. Using additional sequencing data from two closely-related species (*P. campbelli* and *P. roborovskii*), we show that recombination suppression on Xp appears to be independent of major structural rearrangements. The suppressed Xp arm was enriched for several transposable element families and de-enriched for genes primarily expressed in the placenta, but otherwise showed similar gene densities, expression patterns, and rates of molecular evolution when compared to the recombinant Xq arm. *Phodopus* Xp gene content and order was also broadly conserved relative to the more distantly related rat X chromosome. Collectively, these data suggest that widespread suppression of recombination has likely evolved through the transient induction of facultative heterochromatin on the *Phodopus* Xp arm without major changes in chromosome structure or genetic content. Thus, dramatic changes in the recombination landscape have so far had relatively subtle influences on overall patterns of X-linked molecular evolution.

**Significance Statement:** Sex chromosome evolution represents a dynamic process of genomic specialization that is thought to be dependent on evolution of recombination. Here we use genome sequencing and genetic mapping to show that one arm comprising the majority of the X chromosome in a species of dwarf hamster has largely lost the ability to recombine in males and females. Although these dramatic shifts in recombination frequencies might eventually lead to sex chromosome degeneration, loss of recombination on this arm is associated with relatively minor changes in chromosome structure and gene contents in this species. These results underscore the conservation of the X chromosome across mammals, and allow us to test predictions about how genetic recombination influences sex chromosome evolution.

---

The recurrent evolution of sex chromosomes represents one of the most extreme examples of genomic specialization. One hallmark of sex chromosome evolution is that the chromosome present only in the heterogametic sex often evolves a highly reduced, repetitive structure consisting primarily of a sex determination switch and genes involved in reproduction (reviewed in (1–3)). A common outcome of this process is an orthologous but heteromorphic sex chromosome pair with a single, degenerated sex-limited chromosome and a largely intact, recombining chromosome found in both sexes. Heteromorphic sex chromosomes are found in several of the most commonly studied organisms, including XY male heterogametic systems (most mammals and *Drosophila*), and ZW female heterogametic systems (birds and *Lepidoptera*). However, examination of these relatively ancient sex chromosomes reveals only the late stages of a presumably dynamic and complex process of structural evolution, functional specialization, and gene loss (1).

As sex chromosomes have been surveyed in more taxa, it has become clear that the amount of degeneration on the sex-limited chromosome (e.g., the Y or W) can be decoupled from chromosome age (4) and can vary between very closely related species (4–6). Recombination is the primary determinant of this dynamic process. Once a locus with sex-specific effects arises on a chromosome with a sex determination gene, sexually antagonistic selection should favor maintaining linkage between the two loci through suppressed recombination (2). As this process continues, regions with suppressed recombination are expected to lose functional genes and accumulate repetitive elements (7), and ultimately increase the proportion of the chromosome packaged as heterochromatin (8). Chromosomal inversions are thought to be a primary mechanism for recombination suppression, as they impede pairing and synapsis between homologous chromosomal regions in heterozygotes (9) and thereby link sexually-antagonistic alleles to sex determiners (10). Consistent with this model, complex structural rearrangements are a common feature differentiating heteromorphic sex chromosomes. However, inversions are only one possible mechanism for suppressed recombination (11), and it remains unclear if the rapid accumulation of inversions on sex chromosomes are a cause or consequence of broader recombination suppression (3).

In contrast, the X or Z chromosomes usually undergo free recombination in the homogametic sex (e.g., XX females or **A B** ZZ males). As a consequence, shared sex chromosomes often ^3^ maintain some similarity to their inferred ancestral (autosomal) form and tend to be more conserved, but also have the potential to evolve rapidly. For example, mammalian X chromosomes tend to show broad conservation of gene content and order across placental mammals, reflecting a common origin and strong purifying selection related to dosage compensation (12, 13) and likely other functional dynamics (11, 14). At the same time, the X chromosome also shows several signatures of rapid evolution relative to the autosomes, including more rapid protein-coding evolution (i.e., faster-X evolution; reviewed in (15)) and enrichment for genes with both male- and female-specific functions (16, 17). Faster-X evolution is generally thought to reflect accelerated adaptive change due to immediate exposure of recessive beneficial mutations in males (18). However, both the strength of faster-X evolution and the pattern of sex-biased gene content varies considerably across taxa (1), likely reflecting variation in effective population sizes, the degree of female-biased transmission of the X, mechanisms of dosage compensation (19), and epigenetic inactivation during certain developmental processes (e.g., meiotic sex chromosome inactivation in mammalian testis (20)). Understanding how all of these processes shape mammalian X chromosome evolution has been limited by the relatively ancient common origin and general structural conservation shared among taxa with sequenced genomes.

A rich history of mammalian cytogenetic research points to many potential exceptions to a conserved X paradigm that might help advance understanding of sex chromosome evolution (21, 22). One such exception may occur within the Cricetid rodent genus *Phodopus*, comprised of three species: *Phodopus sungorus* (Siberian or Djungarian hamster), it’s closely-related sister taxon *P. campbelli* (Campbell’s dwarf hamster), and *P. roborovskii* (Desert hamster). All three species are endemic to xeric steppe and desert habitats of central Asia (23–25). *Phodopus* sungorus and *P. campelli* have been developed as laboratory models to understand circadian and circannual plasticity in physiology, reproduction, and behavior (e.g., (26–28)). *Phodopus* species are also noteworthy for their unusually large sex chromosomes. The submetacentric *P. sungorus* X chromosome comprises ∼ 10% haploid female karyotype (29), approximately twofold more than the mammalian standard karyotype (30), with the karyotypically shorter (Xp) arm appearing largely as condensed constitutive (C-banding) heterochromatin (29, 31). The heterochromatic Xp arm is presumed to represent a massive expansion of highly repetitive (presumably non-genic) sequences in P*hodopus* (29), with a short distal pseudoautosomal region where all male-specific recombination occurs. Subsequent immunostaining (32) and genetic crossing (33) experiments have confirmed dramatic suppression of female recombination on the presumed Xp arm in *P. sungorus, P. campbelli*, and/or their F1 hybrids. However, the genic content of Xp is unknown and X-linked incompatibilities are also primary determinants of hybrid male sterility (32) and disrupted hybrid placental development (33) in intercrosses between *P. sungorus* and *P. campbelli*, suggesting that functionally-relevant components of the Xp arm may be rapidly evolving.

Here we combine shotgun and long-range proximity ligation sequencing with genetic mapping data to construct the first chromosome-scale *de novo* genome assembly for the Siberian dwarf hamster (*P. sungorus*). We then use multi-tissue transcriptomes, shotgun genome sequencing from *P. campbelli* and *P. roborovskii*, and comparative genome data from other mammals to examine the evolution of chromosome structure, genetic content, and protein-coding sequences across the *Phodopus* X chromosome.

## Results

### *Phodopus* chromosome-scale genome assembly and annotation

We generated an initial *de novo* assembly for a single *P. sungorus* female based on 742 million paired Illumina reads (150 bp PE, Illumina HiSeq X), resulting in near complete assembly of the estimated non-repetitive genome and 91% of the estimated total genome (estimated size 2,310 Megabases or Mb). The total assembly covered 2,113.3 Mb (98,297 contigs, contig N50 = 51.5 kilobases or kb) collected into 69,381 scaffolds (scaffold N50 = 79.7 kb). These data were then augmented with long-range proximity ligation data using complementary Dovetail Chicago and Hi-C methods. The end result was a highly contiguous scaffolded assembly (scaffold N50 = 165.75 Mb; scaffold N90 = 30.61 Mb) with ∼ 90% of the build contained within 14 scaffolds (Table S1); a dramatic improvement relative to a previously published draft Illumina shotgun assembly for this species (contig N50 = 2.2 kb; scaffold N50= 4.8 kb (34)). We then evaluated the quality of the genome assembly by re-mapping the short reads and comparing diploid genotype likelihoods for matched and mismatched bases relative to the haploid assembly of the longest scaffold for each chromosome as implemented with Referee (35). We estimated a per-base error rate of 4.22^10*−*5^, with 67.6% of evaluated bases receiving a quality score over 80 (Fig. S1).

We previously generated a genetic map for *Phodopus* (33) composed of 14 major linkage groups based on a backcross mapping experiment [F1 female (*P. campbelli* female x *P. sungorus* male) x *P. campbelli* male]. Both *P. sungorus* and *P. campbelli* have 13 mostly metacentric autosomes and a sex chromosome pair (2N=28; FN=51 (24, 25)), suggesting recovery of one linkage group per chromosome. To combine our genetic and physical maps, we mapped the ordered markers used to construct the genetic map to the genome assembly and anchored 37 scaffolds (99% of the total genome assembly) into 14 chromosomal linkage groups. Nine of the large scaffolds corresponded to single linkage groups, likely representing eight of the 13 autosomes and the X chromosome. Four autosomes were recovered in two large scaffolds, while chromosome 5 was more fragmented with six assembled scaffolds (Table S2; numbering based on *de novo* genetic map lengths from (33).

Finally, we used available mouse protein data, *Phodopus* transcriptome data sampled from 10 tissues, and *ab initio* predictions to generate and refine gene models. After filtering, we annotated 27,906 protein-coding and non-coding genes, including 87% (264 of 303) of the single-copy orthologs from the set of eukaryotic benchmarking genes (BUSCO, odb9; (36)). Of the annotated genes, 23,736 corresponded to protein-coding transcripts in the house mouse (mm10), of which 14,411 were 1:1 orthologs.

### Widespread suppression of X-linked recombination on the Xp arm

Our previous mapping experiment indicated map compression on one end of the X chromosome (33). To quantify per chromosome recombination rates and the magnitude of X-linked compression, we compared the number of base pairs anchored on each chromosome to its genetic length on the linkage map. Genome build size (Mb) predicted 76.6% of the variance in recombination frequencies (map distance in centimorgans or cM), with 11 chromosomes falling within the 95% confidence interval of the linear model (Fig.1A). The X chromosome was ∼ 1.7x the expected physical length predicted by genetic map distances (71.6 cM expected vs 41.3 cM observed; Fig.1). Recombination rates vary considerably across mammalian chromosomes and species, with X-linked rates typically lower in rodents in sex-averaged studies (37). We estimated an X-linked female recombination rate of 0.359 cM/Mb (Fig.1B), compared to a genome-wide recombination rate of 0.55 cM/Mb which is similar to genome-wide rates in mice (0.523 cM/Mb) and rats (0.551 cM/Mb, rates from (37)).

**Fig. 1.**
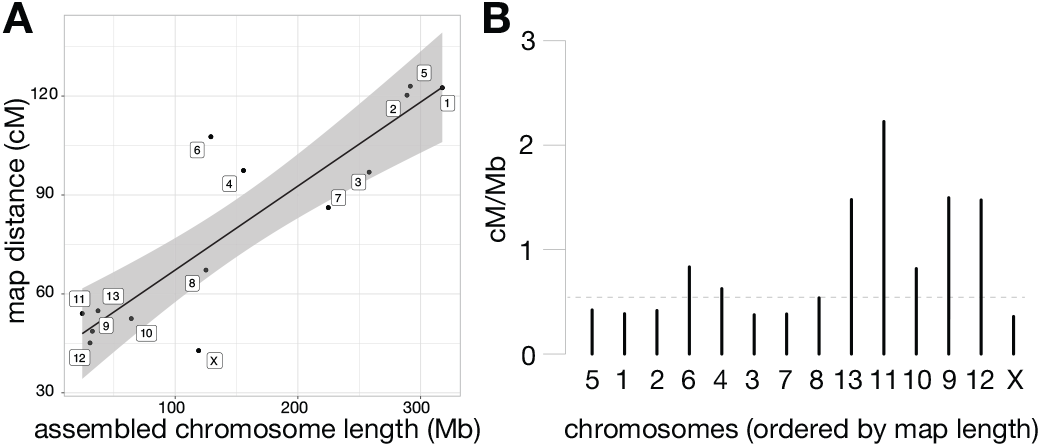
Assembled length vs genetic map distance for all chromosomes. (A) Assembled chromosome length, determined by the total length of scaffolds anchored to the chromosome level, compared to the length of each chromosome in the genetic map. Black line shows the linear relationship between genetic and physical distance, with the gray shading indicating the 95% confidence interval of the linear model. (B) Recombination rate (cM/Mb) for all chromosomes, ordered by genetic length in the anchoring map. Genome-wide recombination rate is indicated with a dashed line. Chromosome numbers are retained from Brekke *et al* (2021), which named linkage groups based on genetic length from a *de novo* map.

Centromeres are dynamic chromosomal features that include diverse families of repeats (38) and can have functional epi-alleles that vary even within populations (39), making it difficult to localize centromeres in genome assemblies. Therefore, we examined crossover events across the assembled chromosomes and used the inherent reduction in recombination that results from condensed pericentric heterochromatin during meiosis to approximate the location of the centromeres in our assembly. Of the 13 autosomes, 10 displayed characteristic sinusoidal patterns with increased recombination near the distal regions of the chromosome arms (Fig. S2), as seen in other mammals with metacentric centromeres (40). Two chromosomes showed patterns consistent with an assembly error (chromosomes 9 and 11), which were localized by the bounds of the markers in the genetic map, but were unable to be further resolved without generating additional higher coverage long-read data for *P. sungorus*. In contrast, the X chromosome showed a partial sinusoidal pattern only for the first third (∼ 42 Mb) of the chromosome, with female recombination showing a reduction at the presumed centromere and then remaining suppressed along the rest of the X (Fig.2). Based on previous observations that recombination is mostly absent from the condensed Xp (32), we used the inflection point of recombination suppression to partition the X chromosome assembly into two arms, ∼ 42 Mb (Xq) and ∼ 77 Mb (Xp) long respectively. Under this model, the Xp arm (i.e., the shorter arm under standard convention) actually comprises 65% of the assembled X *Phodopus* chromosome (77 of 119 Mb).

**Fig. 2.**
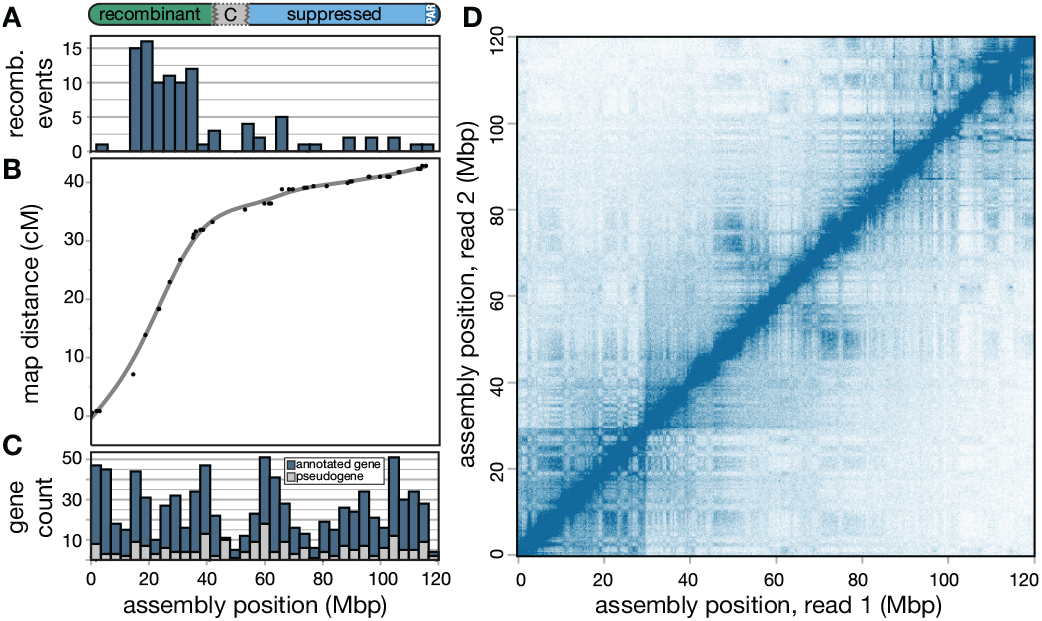
Recombination suppression, gene content, and chromatin configuration of the *Phodopus* X chromosome. (A) Recombination events are predominantly localized to a single arm of the X chromosome, which results in (B) an arm-specific map compression due to suppressed recombination. (C) Genes with complete gene models (blue) and putative pseudogenes with only expressed transcript support (gray) are distributed uniformly across the chromosome, despite the varying recombination levels. (D) HiC chromatin interaction plot, showing short- and long-range interactions between points on the X chromosome with a 250 Kbp resolution and a square root coverage normalization.

In the *P. sungorus* karyotype, the “shorter” Xp arm usually appears as condensed chromatin, which obscures its physical length relative to the less condensed Xq arm. The distal end of the Xp also harbors the pseudoautosomal region (PAR), which is the location of X-Y homology, synapsis, and all X-linked recombination in males. To verify that the longer assembled arm was indeed the Xp, we generated whole genome shotgun (WGS) sequence data for one outcrossed male *P. sungorus* and two reciprocal hybrids between *P. sungorus* and

*P. campbelli*. We then used normalized patterns of female:male sequencing coverage and SNP density to localize the PAR to the distal 3 Mb of the longer arm (Fig. S3). Hereafter, we refer to the suppressed and recombinant regions as the Xp and Xq chromosomal arms, respectively, with the caveat that additional work is needed to verify the exact location of the centromere. The overwhelming majority of observed female crossover events in our mapping panel occurred along the Xq arm, with an elevated estimated recombination rate of 0.78 cM/Mb relative to the *Phodopus* genome-wide rate of 0.55 cM/Mb. Conversely, female recombination was severely suppressed across the entire Xp arm (0.13 cM/Mb), with 74 Mb of 77 Mb presumably experiencing little or no recombination in either sex.

The observation of dramatically reduced recombination across Xp is broadly consistent with heterochromatic suppression of recombination in females (32). However, our recombination data derives from F1 females between *P. sungorus* and *P. campbelli* and, therefore, could also reflect species-specific structural differences. To address this alternative hypothesis, we evaluated short read and long read mapping data, and compared chromatin configuration maps between the species. Comparisons of breakpoints identified by mapping of short read pairs support a ∼ 20 Mb inversion on the distal region of Xp (spanning 95-115 Mp on the *P. sungorus* assembly, Fig. S4), which was further supported by 45 Nanopore long reads spanning the inferred breakpoints. Species differences in chromosome order were also detected from reductions in close chromosomal contacts in the *P. campbelli* HiC data at 115 Mb, compared to the *P. sungorus* assembly (Fig. S5C). This apparent inversion only encompassed 26% of the Xp arm (20 of 77 Mb), and was insufficient to explain the broad recombination suppression extending well outside of this region.

### Conservation of ancestral gene content and order despite the evolution of extensive repressive heterochromatin

Extraordinary conservation of gene orders is one of the hallmarks of mammalian X chromosome evolution (41), and may reflect strong selection preserving three-dimensional chromosomal structure of the X (14). To test whether dwarf hamsters have broadly maintained X chromosome synteny, we aligned the dwarf hamster X chromosome to the rat, mouse, dog, and human reference genomes. We also compared the relative order of our annotated genes to orthologous genes on the rat X chromosome. The distal end of the dwarf hamster Xp showed a similar pattern of rearrangement relative to both the dog and human references (Fig. S6). We detected several mouse-specific rearrangements across the X relative to other species, consistent with previous reports (Gibbs et al 2004; Brashear et al 2021). However, we found high conservation of X-linked gene orders between the *P. sungorus* dwarf hamster reference and rat, with an apparent inversion on the distal end of Xp and slight re-ordering of genes on the distal end of Xq (Fig.2). The inversion breakpoints on Xp match up with the inversion detected between *Phodopus* species (Fig. S4, S5), suggesting that *P. campbelli* retains the same, presumably ancestral, gene order as rat.

Constitutive heterochromatic chromosomal regions are generally assumed to be depauperate of expressed genes, reflecting the repressive effects of condensed chromatin on transcription and enrichment of tandem repeats near pericentromeric and telomeric chromatin (42). Given this and the relatively large size of the *Phodopus* X chromosome, the condensed and darkly staining Xp arm has been assumed to be highly repetitive and largely non-genic (29). Contrary to these predictions, we found that the Xp arm harbors approximately 54% of expressed genes on the *Phodopus* X chromosome (379 of 697 genes, Table S3), which is similar to the proportion of genes found on the orthologous and largely collinear arm in rat (57%, 472 of 824, Table S3). The Xp region did have fewer intact genes than expected given its proportional length in the dwarf hamster assembly (379 genes observed vs 432 genes expected, Fisher’s exact p < 0.0001). However, this pattern was also apparent in the orthologous region of the rat X (Fisher’s exact p < 0.0001), suggesting an ancestral reduction in gene density on Xp rather than gene loss subsequent to recombination suppression.

Even though we found many intact genes on the dwarf hamster Xp, it is possible that the presence of heterochromatin could still impact gene function by broadly suppressing levels of gene expression. When we considered all the tissues except for testis (Fig. S7A), there was a slight reduction in expression on Xq relative to autosomes (pairwise Wilcoxon, p = 0.018) but no difference in the expression levels of Xp relative to the autosomes (pairwise Wilcoxon, p = 0.702) or Xq (pairwise Wilcoxon, p = 0.126). Testis was examined separately (Fig. S7B), as the X chromosome likely undergoes meiotic sex chromosome inactivation (MSCI) and postmeiotic repression during spermatogenesis, which is expected to reduce expression of many X-linked genes. Reductions in testis expression consistent with MSCI were apparent for both Xq (pairwise Wilcoxon, p < 0.0001) and Xp (pairwise Wilcoxon, p < 0.0001), with no difference in expression levels between the arms (pairwise Wilcoxon, p = 0.44).

When we included transcribed pseudogenes in the counts of gene distribution across arms, rat no longer showed a deficit of genes on the region orthologous to hamster Xp (660 of 1084; Fisher’s exact p = 0.1194), indicating more ancient loss of functional genes in this region in rodents that pre-dates recombination suppression in *Phodopus*. When we included putative pseudogenes in dwarf hamster (i.e., expressed genes in reference-guided transcriptomes without a complete gene model in the annotation), *Phodopus* continued to have slightly fewer genes than expected based on length (511 of 896; Fisher’s exact p < 0.0001). This may indicate that dwarf hamsters have lost pseudogene expression relative to the rat, though it is more likely that this difference merely reflects more complete rat annotation.

Finally, if Xp were comprised of constitutive heterochromatin we should detect a clear signal of insulation from within-arm, long-range interactions from our HiC data as is seen in the inactive mouse X (43). In contrast, our HiC results from liver tissue of both *P. sungorus* (Fig.2D) and *P. campbelli* (Fig. S5B) revealed smaller interspersed blocks of topographically-associated domains (TADs) that indicate active transcription (43). Thus, there was relatively little insulation on the arm with suppressed recombination, indicating that the heterochromatic state seen in metaphase karyotypes is tissue-dependent or perhaps transient in at least some transcriptionally active tissues.

### Expansion of transposable element content on Xp

Another potential consequence of reduced recombination is the accumulation of transposable elements (TEs) due to a reduced efficacy of selection (44). To test for an accumulation of transposable elements and other repeats on Xp, we annotated repetitive sequences in our genome build and the rat genome (version rn6; (45)). If suppressed recombination leads to the accumulation of TEs and repeats, we might expect to see a relative enrichment of repeat families on the dwarf hamster Xp relative to the homologous arm in rat. We found support for this hypothesis, as repeat families enriched on Xq were shared between the two species, but repeat families enriched on Xp were found enriched predominantly in hamster only. Dwarf hamster and rat shared patterns of TEs enriched on the female-recombining Xq and several transposable element families (e.g., SINEs and LINEs) were de-enriched on Xp and the homologous arm in rat (Fig.4F-I, Table S4), likely reflecting the shared evolutionary history of this region of the X chromosome. However, three of the four transposable element families enriched on Xp in dwarf hamster showed no evidence for repeat expansions in rat. Only the long terminal repeat (LTR) family ERV1 was also enriched in the orthologous region of the rat X (Fig.4E). Two families (DNA/EnSpm and LTR/MaLR, Fig.4B, D) enriched on Xp in dwarf hamster were significantly de-enriched in rat (Table S4). We also found that the Ngaro family of long terminal repeats (LTRs, Fig.4A) was highly enriched on only the recombination-reduced Xp (170 of 179 copies, Fisher’s exact p < 0.0001). In addition to TEs, we identified enrichment for other repetitive element classes (simple repeats, low complexity sequence, and repeat families of unknown identity, Fig.4C) on the *Phodopus* Xp, but not the recombining orthologous arm in rat. Thus, suppression of recombination appears associated with the accumulation of some repetitive sequence on the *Phodopus* X chromosome.

### Molecular evolution of the X chromosome

To quantify patterns of molecular evolution on and off the X chromosome in dwarf hamsters, we evaluated the ratio of per site nonsynonymous (dN) and synonymous (dS) nucleotide changes in protein coding sequences (dN/dS). We estimated gene-wide rates in two parallel sets of four hamster or mouse species, each spanning similar evolutionary timescales (Fig.5A, B). Consistent with previous results (46, 47), we found faster-X protein-coding evolution for both mice (*Mus* dN/dS_*auto*_ = 0.15; dN/dS_*X*_ = 0.19, Wilcoxon p < 0.0001) and hamsters (*Phodopus* dN/dS_*auto*_ = 0.17; dN/dS_*X*_ = 0.21; Fig.5D, Wilcoxon p < 0.0001). We also found a decrease in the rate of synonymous changes on the X chromosome for both groups (Wilcoxon *Mus* p < 0.0001, *Phodopus* p < 0.0001) without a corresponding reduction in non-synonymous changes (Fig.5C). Interestingly, dS was similar between the Xp and Xq arms (Fig.5E) and we did not observe differences in dN/dS between the two arms (Fig.5F).

**Fig. 3.**
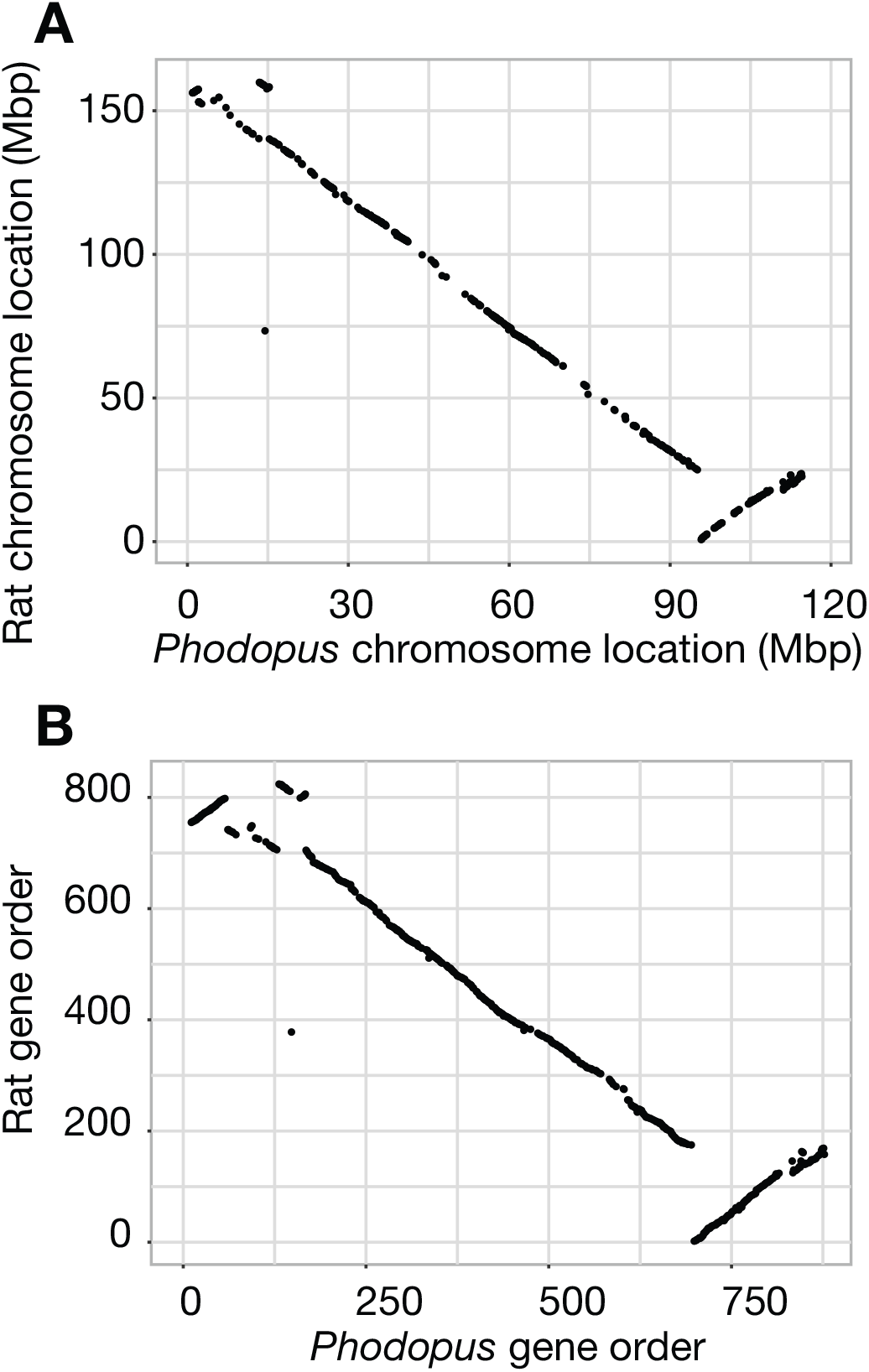
Conservation of chromosome location and gene order between dwarf hamster and mouse. Gene start position (Mb) (A) and relative order (B) are largely conserved between dwarf hamster and rat along the X chromosome.

**Fig. 4.**
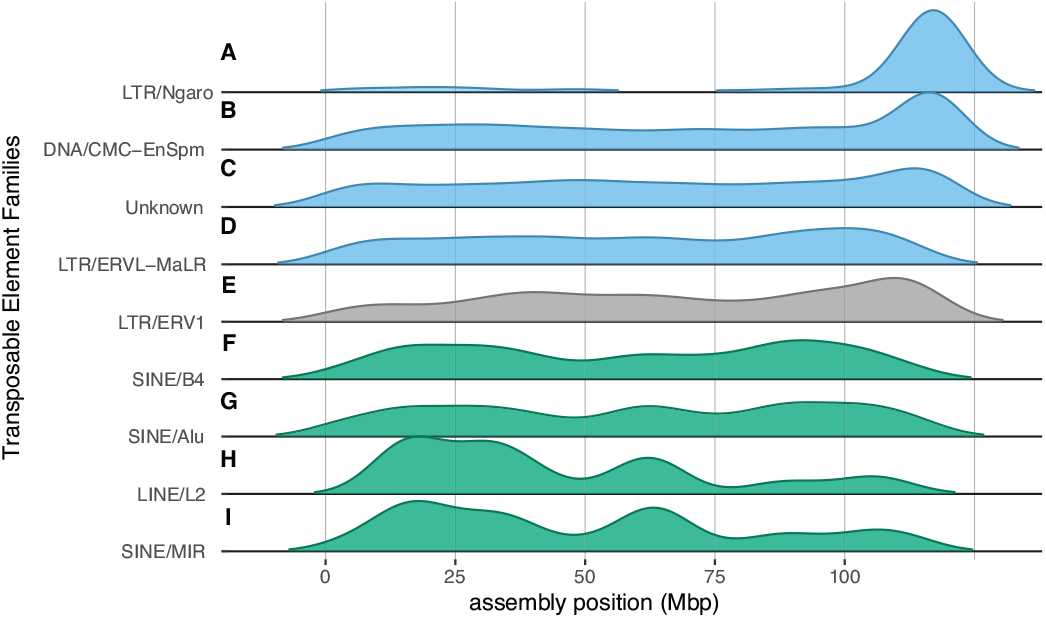
Spatial distributions of the classes transposable elements significantly enriched on an arm of the *Phodopus* X chromosome. Distributions of TE families across the *Phodopus* X assembly, position in Mbp. The top four classes (A-D; blue) are enriched on Xp only in dwarf hamster, ERV1 (E, gray) is enriched on Xp in both dwarf hamster and rat, and the bottom four classes (F-I; green) are enriched on Xq in both dwarf hamster and rat. Significance determined by binomial test, see Table S4 for values.

**Fig. 5.**
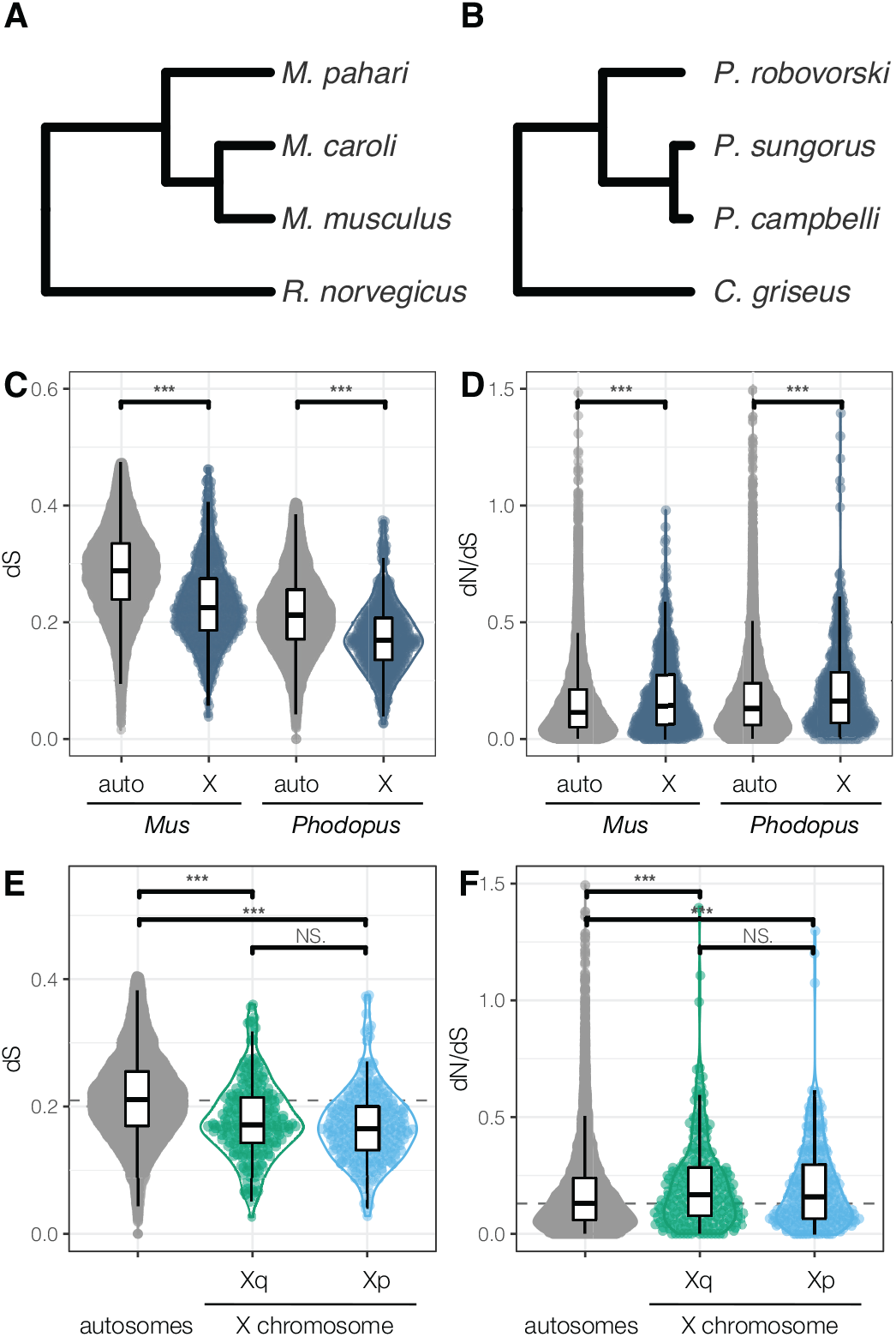
Evolutionary rates on autosomes and X chromosome arms in Phodopus. Gene-specific rates of change in synonymous nucleotides (dS, C and E), and the ratio of non-synonymous to synonymous rates of change (dN/dS, D and F) for coding genes on the autosomes and X chromosome arms, calculated separately for Mus (A) and Phodopus (B). For Phodopus, we also compared rates between the recombining and non-recombining arms of the X chromosome (E, F). Letters indicate significant differences within panel at p < 0.05, Wilcoxon nonparametric multiple comparisons test. Dashed lines indicate genome-wide median.

To test whether dwarf hamsters display sex-biased gene expression on the X chromosome, we examined gene expression across nine tissues (brain, heart, kidney, liver, muscle, spleen, uterus, placenta, and testis) using a specificity index (*τ*) to identify genes with tissue-enriched expression (Table S5; Fig.6). Approximately the same number of genes were expressed in each tissue (average = 17,904 ± 542 genes, range 15,117-19,693 genes), and the majority of these genes were annotated as protein-coding based on orthology with *Mus* (93%, 23,736 of 25,506 genes) The testis was enriched for tissue-specific expression; nearly one-third of all genes with a tissue specificity index (*τ*) greater than 0.8 were expressed in the testis (2,918 of 8,895 genes, Table S5), consistent with previous studies (48). We also detected enrichment on the X chromosome for testis-specific genes (164 genes, hypergeometric p < 0.0001, Fig.6A) and placenta-specific genes (50, hypergeometric p < 0.0001, Fig.6B). When considered separately, each arm of the X chromosome was enriched for testis-specific expression (Xq, 67 genes, hypergeometric p = 0.0005; Xp, 97 genes, hypergeometric p < 0.0001, Fig.6A). However, only the female-recombining Xq arm was enriched for placenta-specific genes (29 genes, hypergeometric p < 0.0001, Fig.6B).

**Fig. 6.**
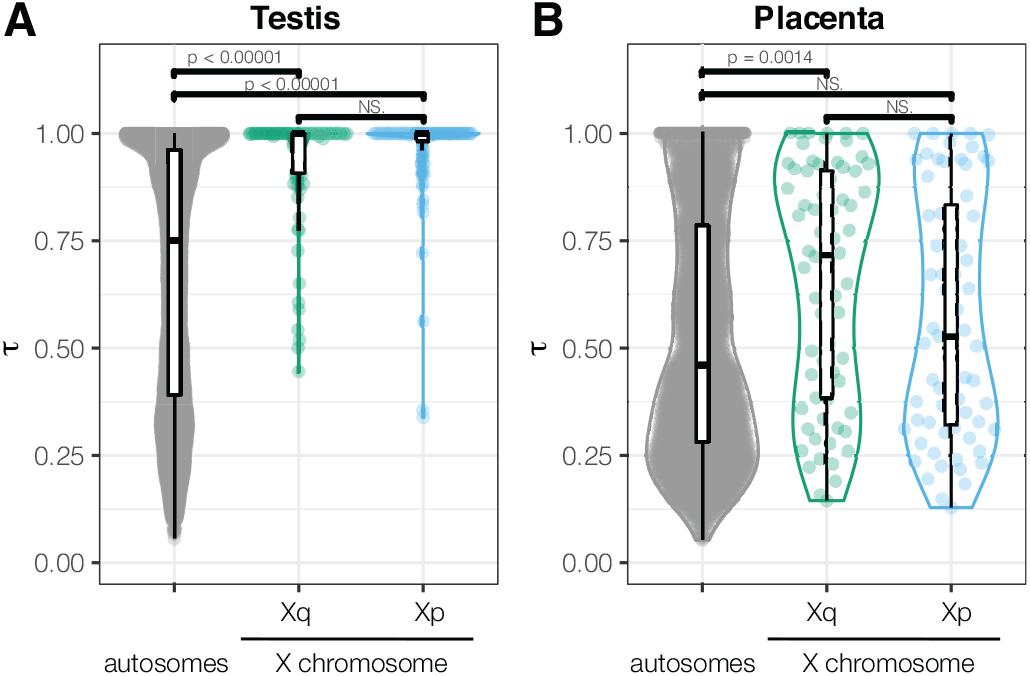
Tissue specificity score *τ* for genes with highest expression in testis (A) and placenta (B), by X chromosome region. Higher values of *τ* indicate higher tissue specificity in expression pattern.

When considering all genes expressed in the testis, rather than those with testis-specific expression, we no longer found an enrichment on the X chromosome (447 observed, 489 expected; Fisher’s exact p = 0.0572). The X showed a general de-enrichment of all placenta expressed genes (375 observed, 445 expected; Fisher’s exact p = 0.0037), which was likely driven by a greater reduction of expressed genes on Xp (199 observed, 241 expected; Fisher’s exact p = 0.0048) compared to Xq (186 observed, 202 expected; Fisher’s exact p = 0.1016).

We also evaluated protein-coding sequence evolution in the four tissues with at least ten tissue-specific genes on each arm of the hamster X chromosome (brain, testis, placenta, and uterus, Fig. S8). In *Mus*, previous work has shown increased dN/dS in male-biased genes on both the X and the autosomes, but no evidence for faster evolution in female-biased genes (46). In hamsters, we detected faster evolution of testis-biased genes across the autosomes and X compared to genome-wide values (Wilcoxon p <0.0001), with dN/dS for these genes on Xp slightly elevated relative to testis-biased autosomal genes (p = 0.037, Fig. S8B). We also found elevated dN/dS for X-linked placenta-specific genes, with placenta-biased genes evolving faster on Xq (Wilcoxon p = 0.012, Fig. S8C). Tissue-specific genes are often associated with elevated dN/dS relative to the genome-wide baselines (49), presumably reflecting reduced evolutionary constraints. In this admittedly limited survey, we detected significantly reduced rates for autosomal brain genes (Wilcoxon p < 0.0001), suggesting that tissue-specificity *per se* does not account for increased rates of X-linked evolution in dwarf hamsters.

## Discussion

Mammalian X chromosomes are often characterized by specialized gene contents that also tend to be highly conserved between species, reflecting the late stages of a presumably dynamic process shaped by diverse selective pressures. Here we show that recombination suppression has evolved along most of an entire arm (∼ 65%) of the X chromosome in *Phodopus* dwarf hamsters. These findings provide an opportunity to evaluate the causes and evolutionary consequences of widespread recombination suppression on a mammalian sex chromosome.

### The evolution of recombination suppression on the *Phodopus* X chromosome

Most studies that have examined recombination suppression in natural systems have focused on segregating structural inversions, which prevent the inverted region from forming successful Holliday junctions between alternate haplotypes. Pairing of sequence between sister chromatids is critical to successful crossing over during homologous recombination (50). On autosomes, segregating inversions have been found to prevent recombination between adaptive loci co-localized in the genome, linking together sets of genes for local adaptation (i.e., ‘supergenes’ in birds, butterflies, and Mimulus; reviewed in (51)). Inversions are also generally assumed to play a critical role in the evolution of heteromorphic sex chromosomes (11, 52), including stratified divergence between the mammalian X and Y chromosomes through a series of rearrangements (53). Indeed, inversions have been shown to maintain a heritable unit consisting of sexually-antagonistic alleles and a sex determination gene in young, homomorphic sex chromosomes in cichlid fishes (10). New inversions arise even on relatively old and conserved sex chromosomes—the mammalian X and avian Z chromosomes are enriched for inversions relative to autosomes (54). Theory predicts that the higher efficacy of selection on the X should limit the accumulation of deleterious mutations in inversions in the short term, which ultimately should facilitate higher inversion fixation rates in the long term when compared to autosomes (54).

Despite the intuitive link between structural changes and reduced recombination, we did not find large chromosomal rearrangements that easily explain the widespread suppression of recombination observed on the dwarf hamster Xp. Although our recombination map was generated with inter-species hybrids (33), the lack of support for an inversion model is unsurprising in this instance. The Xp usually appears to be highly condensed within *Phodopus* species, with clear evidence for suppression of female recombination on the presumed Xp arm within both *P. sungorus* and *P. campbelli* based on immunolocalization of the MLH1 mismatch repair protein (32). Thus, there are multiple lines of evidence for recombination suppression in crosses both with and between *Phodopus* species.

Karyotypes in *P. sungorus* and *P. campbelli* show strong C-banding of the Xp arm (29, 31), a hallmark of constitutive heterochromatin. However, these early studies also noted some variation between individuals in the condensation of Xp in different chromosome spreads, suggesting that the “constitutive” heterochromatin in this region was unlike the condensed, gene-poor regions of chromosomes found predominantly near centromeres and telomeres. Our findings confirm that the heterochromatic state of Xp is likely transient rather than classically constitutive. *Phodopus* Xp shows a chromatin state consistent with that of a transcriptionally active chromosome in liver tissue (Fig.2; Fig.S5), and the arm also has many genes expressed across tissues with no broad reduction in gene expression (Fig S7). One possible explanation is that Xp is condensed using canonical constitutive chromatin mechanisms, but in an inconsistent manner. Pericentromeric constitutive heterochromatin has been historically viewed as unchanging in the face of tissue type and environmental factors. However, research has increasingly uncovered dynamic complexity in the activity of the constitutive (H3K9me3) histone modification machinery (42), including expression of pericentromeric repeats during development and in actively proliferating tissues (55). The *Phodopus* X chromosome appears to have maintained broad conservation of gene order over the last ∼ 23 million years since it shared a common ancestor with murine rodents (Fig.3, (56)). Therefore, many interspersed repeats associated with constitutive chromatin would presumably have had to have evolved along the Xp without major changes in synteny.

Perhaps more likely, condensation of Xp may be occurring via facultative heterochromatin pathways, such as that used in X chromosome inactivation (XCI) in females (57). These mechanisms are often context-dependent (e.g., sensitive to developmental timing and environmental cues) in addition to being sequence-dependent. XCI is initiated at the X inactivation center, where long noncoding RNA (lncRNA) *Xist* coats the X chromosome that will undergo inactivation (58). *Xist* then binds intermediary proteins to recruit the repressive Polycomb complex, a structure that mediates transcriptional silencing genome-wide (59). In this way, the universal machinery for the formation of facultative heterochromatin is recruited in an X-specific manner as XCI progresses. The X chromosome is prepared for heterochromatin formation even before *Xist* marks the inactive X; the histone deacetylase HDAC3 is pre-loaded at enhancers along the X chromosome and plays a critical role in initiating chromosome condensation (57). Thus, the X chromosome has pre-existing sequences across its length that prime the formation of condensed heterochromatin in females. It is plausible such mechanisms could have been co-opted or expanded from the standard XCI pathway, inducing the transient heterochromatin state observed on Xp. Future studies examining histone marks indicative of facultative and constitutive heterochromatin are needed to test the epigenetic mechanisms underlying the formation of transient heterochromatin on Xp.

Considerable literature has focused on the role that inversions play in generating recombination suppression (e.g., (52, 60, 61)). However, there remains relatively little direct evidence that rearrangements act as the primary proximate mechanism underlying the suppression of recombination on sex chromosomes (11).). Regardless of the precise mechanisms, our results point to the relatively recent evolution of epigenetic suppression of recombination across a large portion of a mammalian X chromosome. It is also possible that other sequence changes, such as smaller inversions, nucleotide divergence (62), or TE accumulation (11), generally contribute to reduced X-linked recombination frequencies between *P. sungorus* and P. cambelli. However, such mechanisms cannot not explain suppression within these species (32). In other eukaryotic taxa, reports suggesting that non-inversion mechanisms may be at play in suppressed recombination have been more limited. Sylvioidea songbirds have a ‘neo Z’ region (i.e., an autosomal fusion on the Z chromosome) with a history of complex, progressive recombination suppression and repeat accumulation (63). Perhaps more relevant, recombination suppression in a group of Ascomycete fungi pre-dates inversions between species (64), with a proposed mechanism involving increased DNA methylation generated via the fungus-specific repeat-induced point (RIP) mutation repair pathway (65).

### Evolutionary consequences of recombination suppression

In principle, a long-term reduction of recombination on Xp should lead to different evolutionary trajectories for the two arms of the *Phodopus* X, and provide insights into general models of sex chromosome differentiation. First, the accumulation of repetitive elements and structural variants (i.e., inversions, insertion-deletions (3)) are common features of sex chromosome evolution. The accumulation of sex-linked repetitive elements is generally thought to result from recombination suppression (7, 66) but see (11). Assuming our inference of a non-structural mechanism of suppression is correct, the enrichment for TEs on Xp between *P. campbelli* and *P. sungorus* is likely a consequence of a change in the recombination landscape. Some genomic sequences are predisposed to TE insertions (i.e., TE hotspots (67). However, we observed expansions of several TE families on the *Phodopus* Xp relative to the largely syntenic rat X chromosome, suggesting the differential accumulation of lineage-specific repeats independent of the ancestral sequence landscape of the rodent X chromosome.

The *Phodopus* system may also inform how recombination influences the dynamics of molecular evolution of the X chromosome. One outstanding question about the generation and maintenance of genetic diversity is whether recombination itself is a mutagenic process. This question has been difficult to answer because recombination rate is correlated with other features such as centromere location (40), gene density, and hotspot activity (68). In humans, sequencing evidence from sperm (69) and parent-offspring trios (70) demonstrates that recombination leads to mutation. However, it is less clear whether this phenomenon leads to sequence divergence across taxa (71). Our finding of similar levels of synonymous sequence divergence on the Xp and Xq arms suggests that changes in recombination frequencies have so far been a negligible contributor to substitution rates since the origin of Xp suppression in *Phodopus*.

The mammalian X chromosome generally shows lower synonymous substitution rates, presumably due to higher mutation rates in males (i.e., male-driven molecular evolution; (72) and female-biased transmission of the X chromosome (73). Despite a reduction in baseline mutation, theory predicts more efficient selection in males on the hemizygous X chromosome (18, 74), which may increase the fixation rate for beneficial alleles (i.e., the faster-X effect), especially in the testis and other male-specific tissues. Consistent with this, the X chromosomes of several mammal species show relatively higher rates of protein-coding evolution (dN/dS), which likely reflects more effective selection for recessive beneficial mutations (reviewed in (15)). Our finding of elevated X-linked dN/dS in hamsters parallels results in mice and rats (Fig.5) and indicates that faster-X evolution is likely a general feature of rodent genomes. All else being equal, reduced recombination could limit the efficacy of natural selection on the Xp (i.e., Hill-Robertson interference (75)), which could lead to different rates of protein-coding evolution between Xp and Xq. We did not detect such differences, with both arms showing similarly elevated dN/dS relative to the autosomes (Fig.5).

The X chromosome is also predicted to accumulate genes with sexually antagonistic variation (74), which can lead to an enrichment of sex-biased genes. Similar to predictions for faster-X evolution, the outcomes of selection for sex-biased gene contents are predicted to depend on the dominance of sexually antagonistic mutations. Male-biased genes are predicted to accumulate on the X if male-beneficial mutations are on average recessive. Likewise, given female-biased transmission, female-biased genes are predicted to accumulate on the X if female-beneficial mutations are on average dominant. Interestingly, these theoretical predictions play out differently in different systems (76). In *Drosophila*, the X appears to be strongly de-enriched for male-biased genes (77, 78), while mammals have been reported to be enriched for both male- and female-biased genes with some variation across studies (20, 79). In hamsters, we observed enrichment for both testis- and placenta-specific genes, but not for other sex-specific tissues (i.e., uterus and ovary). These potentially conflicting patterns may be explained by inactivation of the paternal X chromosome in the placenta and other extraembryonic tissues of rodents (80, 81) resulting in predominantly maternal expression of X-linked placental genes. Thus, placental genes are functionally hemizygous, similar to X-linked genes in males. Notably, we found that testis-specific genes were enriched on both arms of the X, while enrichment of placental-specific genes was restricted to the female-recombining Xq arm. These patterns are based on small numbers of genes, but suggest that higher recombination may enhance the efficacy of sexually antagonistic selection in females.

Finally, these unusual evolutionary dynamics on the *Phodopus* X chromosome may also be influencing the rapid evolution of reproductive barriers between dwarf hamster species. Male hybrid sterility between female *P. campbelli* and male *P. sungorus* has been attributed to asynapsis of the sex chromosomes during meiosis (32), indicating that pairing may be inhibited by sequence or structural divergence that has accumulated on the PAR-containing Xp. Crosses between *P. sungorus* and *P. campbelli*) result in extreme hybrid placental overgrowth caused by a large-effect QTL near the P. sungorus Xq-Xp boundary (Fig. S9) that disrupts a network of autosomal genes expressed in the placenta (33). Understanding if and how the evolution of widespread recombination suppression is related to speciation awaits a more detailed dissection of the genetic and mechanistic bases of these hybrid incompatibilities.

## Materials and Methods

### Animal sampling

A single *P. sungorus* female was chosen for *de novo* genome sequencing and assembly from a breeding colony at the University of Montana (UM), originally established in 2011 descending from wild-caught animals collected 1981-1990 (82); see (83) for further details on *P. sungorus* and *P. campbelli* resources). The UM colony was maintained to maximize outbreeding, but levels of inbreeding are nonetheless elevated in these lines (Brekke et al. 2018). In addition to collecting liver from the reference individual for DNA sequencing, we collected seven snap-frozen tissues for use in transcriptome-based genome annotation (brain, heart, kidney, liver, muscle, spleen, and uterus). Testes from a single *P. sungorus* male from the same UM line were also collected to expand the annotation tissue panel. Augmenting the annotation panel, we collected and snap froze skin biopsies from molting *P. sungorus* females as part of an ongoing study on gene expression during seasonal molts. We sampled liver tissue from three additional males for shotgun genome sequencing: one outbred male *P. sungorus* was derived from a cross between a male from our original UM colonies and a female from supplemental stock kindly provided by Matthew Paul (University of Buffalo) in 2019, and two hybrid male embryos from reciprocal F1 crosses between *P. sungorus* and *P. campbelli*. Finally, we sequenced liver tissue from one female *P. campbelli*, derived from the UM research colony, and one male *P. roborovskii* sourced from the pet trade. All animal care and breeding was performed at UM with oversight of the University of Montana Institutional Animal Care and Use Committee (animal use protocols 039-13JGDBS-090413, 050-16JGDBS-082316 and 035-19JGDBS-062519).

### Genome and transcriptome sequencing

Liver tissue was sent to Dovetail Genomics (Santa Cruz, CA) for generation of all sequencing data used in the *de novo* assembly, including Illumina whole genome shotgun sequencing (150 bp PE, Illumina HiSeq X) augmented with Dovetail’s complementary Chicago (relying on artificial chromatin constructed *in vitro*) and Hi-C (relying on intact chromosomes *in situ*) long-range proximity ligation data. The libraries were prepared using Dovetail kits according to the manufacturer’s protocol. For Hi-C libraries, chromatin was fixed in place with formaldehyde in the nucleus and then extracted. Fixed chromatin was digested with *DpnII*, the 5’ overhangs filled with biotinylated nucleotides, and free blunt ends ligated. After ligation, cross-linking was reversed and DNA was purified. Following purification, biotin not contained in the center of the ligated fragment was removed. Chicago libraries were prepared similarly, except DNA was removed from the nucleus and formed into artificial chromatin rather than fixed *in situ*. For all libraries, DNA was sheared to ∼ 350 bp mean size and libraries were generated using Illumina-compatible adaptors.

We also generated an additional Hi-C library from *P. campbelli* liver tissue, processed following the same Dovetail protocols as the libraries generated for genome assembly. Whole genome sequencing libraries from *P. campbelli, P. roborovskii*, and three hybrid males were constructed using the NEBNext Ultra II DNA kit (New England Biolabs, E7645S). RNAseq libraries were constructed from all reference tissues at the University of Montana using an Illumina TruSeq Stranded mRNA library kit (Illumina, 20020595), following manufacturer’s instructions, with the exception of the testis library, which was constructed at Novogene (Davis, CA). All tissues were sequenced on 3 lanes of 150 bp PE Illumina Hiseq 4000 (reference transcriptomes) and 3 lanes of 150 bp PE Illumina Hiseq X (HiC and WGS libraries) at Novogene.

High molecular weight DNA for long read sequencing was extracted from the sequenced *P. sungorus* and *P. campbelli* females using the MagAttract HMW DNA kit (Qiagen, 67563) and assessed using a genomic DNA ScreenTape (Agilent 5067-5365) on a 2200 TapeStation (Agilent, G2965AA). Libraries were constructed and sequenced in the UM Genomics Core (Missoula, MT) using the MinION flowcell platform (Oxford Nanopore, sequencing kit SQK-LSK109 and flow cell FLO-MIN106D).

### Genome assembly and annotation

An initial shotgun sequence-based assembly was generated with 150 bp PE Illumina Hiseq data. Reads were trimmed with Trimmomatic (v 0.39, ILLUMINA-CLIP to remove adaptors, LEADING:20, SLIDINGWINDOW:13:20, MINLEN:23). The hybrid read-based/k-mer assembler Meraculous was used to generate a *de novo* assembly, with a constrained heterozygous (II) model with 79-mers and homozygous peak depth 40.0 selected for depth of coverage and balance between repetitive and heterozygous fractions (84). Reads from the additional Chicago and Hi-C libraries were used in conjunction with the *de novo* assembly as input for Dovetail’s HiRise genome assembly pipeline (85). The first assembly iteration aligned Chicago library sequences to the draft assembly with a modified SNAP mapper (86). HiRise analyzed the distance between Chicago read pairs within scaffolds using a likelihood model, which was used to identify joins and misjoins for scaffolding. The second assembly iteration used the same strategy, scaffolding with the proximity-ligation HiC data.

To assess assembly quality on a per-site basis, we re-mapped the raw reads that went into the assembly and ran Referee (Thomas and Hahn, 2019) on the resulting BAM file to calculate genotype likelihoods while accounting for mapping quality. Sites in which the called base in the assembly does not match one of the two alleles in the most likely genotype are inferred as errors and can be corrected by inserting one of the alleles from that genotype.

We generated a reference-guided genetic map using data from our previously published *Phodopus* genetic map based on high-throughput genotypes (ddRAD tags) from 189 backcross individuals [F1 female (*P. campbelli* female x. *P. sungorus* male) x *P. campbelli* male; (33)]. Processed RAD tags were aligned to the reference genome using BWA (Li and Durban 2009), and variants were called using HaplotypeCaller (gatk, v4; (87)). The genetic map was constructed using rQTL as described previously (33), with the exception that duplicate markers were only dropped from the analysis if they were mapped to the same genomic scaffold. While these markers were redundant for identifying recombination events in the genetic map, they were informative for linking unanchored scaffolds and retained for scaffold placement.

We used the MAKER3 pipeline (v3.01.03; (88)) to annotate gene models using *ab initio* and transcript-based evidence. Initial reference-guided *de novo* transcriptome assemblies were built with the Tuxedo pipeline (bowtie v2.3.4.3; (89); tophat v2.1.1 and cufflinks v2.2.1; (90)) including data from all reference tissues and published data previously used for *de novo P. sungorus* placental transcriptomes (80). These transcriptomes were given to the MAKER pipeline, which combined the *Phodopus* expression data, SNAP and Augustus *ab initio* gene prediction models, and protein homology to *Mus* to predict the location of genes and genic features. These genes were then matched with *Mus* (mm10) and *Rattus* (rnor6) identity through reciprocal best hits BLAST searches (91), using a max e-value cutoff of 1e-3 and then taking the longest remaining match between the two species.

### Identification of the PAR and X-linked structural variation

For each all WGS samples, reads were trimmed (Trimmomatic v0.39; (92)) and aligned to the *P. sungorus* assembly using BWAmem (v0.7.17; (93)). Read depth was called using ‘*samtools depth*’ and averaged across 2500 bp sliding windows (94). For the X chromosome, average coverage in each window was normalized by median coverage, and adjusted to reflect ploidy (ie. female ploidy is 2N). Next, we filtered out all windows that included bases overlapping with any annotated repeat (Repeatmasker (95)). We excluded regions annotated for STRs/low complexity sequences and all windows where the *P. sungorus* female had coverage over the 95% quantile of coverage (1.68X the median) in order to remove artifacts resulting from collapsed repeats or paralogous genes in the assembly. We used the same custom library to identify repetitive regions in both *Phodopus* and *Rattus norvegicus* (rnor6) as was used to repeat mask the genome for annotation.

We used four approaches to evaluate structural variation on the *Phodopus* X chromosome. First, we identified putative interspecific inversions using mapping discordance between *P. campbelli* and *P. sungorus* Illumina paired-end reads mapped to the X chromosome assembly (Breakdancer v1.3.6; (96)). We used analysis based on *P. sungorus* reads to control for any signal due to genome assembly error or repetitive content, and evaluated the difference between the reference and *P. campbelli* reads to find species-specific inversions by normalizing all counts the reference female, which was sequenced to higher sequence coverage. Second, we mapped Oxford Nanopore ultralong reads from P. sungorus and P. campbelli to the assembly using minimap2 (-ax map-ont; v2.17; (97)) and then used IGV (v2.7.2; (98)) to visually verify the large distal inversion by identifying that mapped across the inversion breakpoint, comparing these to a list of 45 Nanopore reads with regions mapped to either side of the inversion. Third, raw HiC reads from both *P. sungorus* and *P. campbelli* were processed with the Juicer pipeline to align and call HiC contact points (99), normalized using the Knight-Ruiz method. These matrices were then visualized and compared using Juicebox (v1.11.08 (100)). Fourth, we evaluated gene order/synteny across mammals. To assess synteny, we downloaded X chromosome assemblies from Ensembl (release 100) for *Rattus norvegicus* (rat, rnor6), *Homo sapiens* (human, GrCh38), and *Canis familiaris* (domestic dog, CanFam3.1). For each species, we used minimap2 (97) to align conserved segments from the *P. sungorus* X chromosome to the target reference (-asm20). Gene order was also evaluated by comparing the locations of *Phodopus* genes with those of the rat genes identified through reciprocal BLAST (described above). Recombination rate was evaluated using the RAD markers used to anchor the genome to the genetic map, using only the sequence length between markers to avoid including sequence without sampled recombination data.

### Gene expression

We performed gene expression analysis using Hisat2 (101) to map reads to the genome assembly, and then used the MAKER-generated annotation to count reads with FeatureCounts (102). We allowed for multiple mapping but proportionally distributed multiply mapped reads across target genes. Counts were normalized using transcripts per million (TPM), and log2 transformed for analysis. All placenta samples were quantified following reference alignment, and the average TPM across samples was used to evaluate tissue specificity. We calculated the tissue specificity metric *τ* as described, using the established cutoff of *τ* 0.8 (48). Expressed genes were also evaluated for overlap protein-coding molecular evolution metrics (see below).

### Molecular evolution

We downloaded orthologous protein coding sequences for five species from Ensembl (release 100): *R. norvegicus, M. musculus, M. caroli, M. pahari*, and *C. griseus*. We used these sequences in conjunction with the *P. sungorus* genome and reference-guided assemblies of *P. campbelli* and *P. roborovskii*, constructed using iterative mapping to the *P. sungorus* to reduce reference bias. These iterative assemblies were as implemented with a modified version of pseudo-it (v.beta 3.1; (103)) that incorporates insertion-deletion variation. For each species, we ran pseudo-it for four iterations and used the UCSC liftOver tool (University of California, Santa Cruz Genomics Institute) to transfer *P. sungorus* gene annotations to each pseudo-assembly.

We split the eight species into two groups of four species that spanned approximate similar evolutionary timescales. For the hamster group (Cricitidae: *P. sungorus, P. campbelli, P. roborovskii*, and *C. griseus*), we used BLAST (91) to obtain orthologous protein coding transcripts between the C. griseus genome and *P. sungorus* by setting a max e-value cutoff of 1e-3 and then taking the longest remaining match between the two species. This resulted in 16,569 single-copy orthologs between these five species. For the mouse/rat group (Muridae: *M. musculus, M. caroli, M. pahari*, and *R. norvegicus*), we used Ensembl annotation to identify 15,772 single-copy orthologs. We aligned orthologous coding-sequences in both groups using MACSE (104), which produces in-frame codon alignments while accounting for frameshifts. First, sequences that contain non-homologous sequences were trimmed with the trimNonHomologousFragments sub-program. In some cases, this removed an entire sequence within the orthogroup, and this entire group was removed from subsequent steps. In total, 2,498 genes were removed from the hamster group and 266 were removed from the mouse group. Next, we aligned sequences with the main MACSE program, *alignSequences* (104). We followed this with a post-alignment trimming step using the trimAlignment sub-program, after which we calculated summary statistics for each alignment and removed those that were short (<100bp), contained a high proportion of sites with gaps or missing data (>20%), contained identical sequences, or contained premature stop codons. In total, using these filters we removed 527 genes from the hamsters and 79 genes from the mice. Following all alignment, trimming and filtering steps, we recovered 13,546 in-frame codon alignments for the hamsters and 15,429 from the mice.

Using these alignments, for both groups, we estimated maximum-likelihood gene trees using IQ-Tree (105) and estimated dN and dS for each gene using the corresponding gene trees in order to minimize the potential effects of discordance (106). Species level phylogenies were also verified from maximum likelihood analysis of a concatenated alignment (Fig.5A, B), but were not used for the analysis of molecular evolution. We used HyPhy (107) to estimate dN/dS using the FitMG94.bf model (https://github.com/veg/hyphy-analyses), estimating a single dN/dS value per gene and summarizing dN and dS separately by summing their values over all branches (Fig.55). For each dataset, we flagged genes with dS values above the top 98th percentile of all genes as having potentially poor alignments and filtered them from subsequent analysis (Fig. S10; 271 Cricitidae genes and 309 Muridae genes).

### Statistical analysis

All basic statistical tests were performed in R v 4.1.0 (108). Non-parametric models were performed with the Kruskal Wallace test function (*kruskal.test*) and posthoc p-values were evaluated with *pairwise.wilcox.test*. Linear models we performed using lm function for standard models, and lmer for mixed models. Enrichment was tested either with hypergeometric test (for enrichment of categories, *phyper*) or Fisher’s exact binomial test (for enrichment based on expected distributions from length, *binom.test*). Multiple testing was corrected for used the Bonferroni Holm method.

## ACKNOWLEDGMENTS

We thank the Good lab for helpful comments on data analysis and interpretation. Ned Place, Matthew Paul, and Robert Johnston provided animal stocks. Kelly Carrick, Jess Wexler, and the University of Montana LAR staff assisted with animal care. Mark Daly, Thomas Swale, and Dovetail Genomics LLC performed genome sequencing and assembly. Denghui David Xing performed Nanopore sequencing at the UM Genomics Core. This research was supported by grants from the Eunice Kennedy Shriver National Institute of Child Health and Human Development of the National Institutes of Health (R01-HD094787 to JMG), a Matching Funds Award from Dovetail Genomics, and the National Science Foundation Graduate Research Fellowship Program (EEKK: DGE-1313190, and KEH DGE-2034612). This study included research conducted in the University of Montana Genomics Core, supported by a grant from the M. J. Murdock Charitable Trust (to JMG). Computational resources and support from the University of Montana’s Griz Shared Computing Cluster (GSCC), supported by grants from the Nation Science Foundation (CC-2018112 and OAC-1925267, JMG co-PI), contributed to this research.

---

**Table S1.** Assembly statistics for Dovetail genome

|  | **Short-read assembly** | **Final Assembly** |
| --- | --- | --- |
| Total Length (Mbp) | 2,113.27 | 2,119.20 |
| L50 (scaffold count) | 7016 | 5 |
| N50 (Mbp) | 0.08 | 165.747 |
| L90 (scaffold count) | 30,428 | 14 |
| N90 (Mbp) | 0.015 | 30.611 |
| Busco (Eukaryota) |  |  |
| Single copy | 246 | 263 |
| Duplicated | 7 | 7 |
| Fragmented | 25 | 9 |
| Missing | 25 | 24 |
| Total | 303 | 303 |

**Table S2.** Chromosome-level anchoring of genome using *Phodopus* genetic map

| **Chromosome** | **Mbp** | **cM** | **Marker**<br>**count** | **All scaffolds** | **Scaffolds > 1Mbp** |
| --- | --- | --- | --- | --- | --- |
| 1 | 317.65 | 122.51 | 208 | 3 | 2 |
| 2 | 288.91 | 120.25 | 174 | 4 | 1 |
| 3 | 258.08 | 96.92 | 148 | 3 | 2 |
| 4 | 155.67 | 97.42 | 137 | 2 | 1 |
| 5 | 291.62 | 122.99 | 134 | 6 | 5 |
| 6 | 129.1 | 107.65 | 135 | 4 | 1 |
| 7 | 224.74 | 86.16 | 118 | 1 | 1 |
| 8 | 125.09 | 67.23 | 74 | 2 | 1 |
| 9 | 32.54 | 48.68 | 68 | 3 | 1 |
| 10 | 64.22 | 52.51 | 52 | 2 | 2 |
| 11 | 37.12 | 54.88 | 46 | 2 | 2 |
| 12 | 24.28 | 54.05 | 47 | 2 | 1 |
| 13 | 30.61 | 45.14 | 42 | 1 | 1 |
| X | 119.12 | 42.79 | 42 | 2 | 1 |
| unplaced | 20.41 | NA | NA | 7719 | 0 |

**Table S3:**
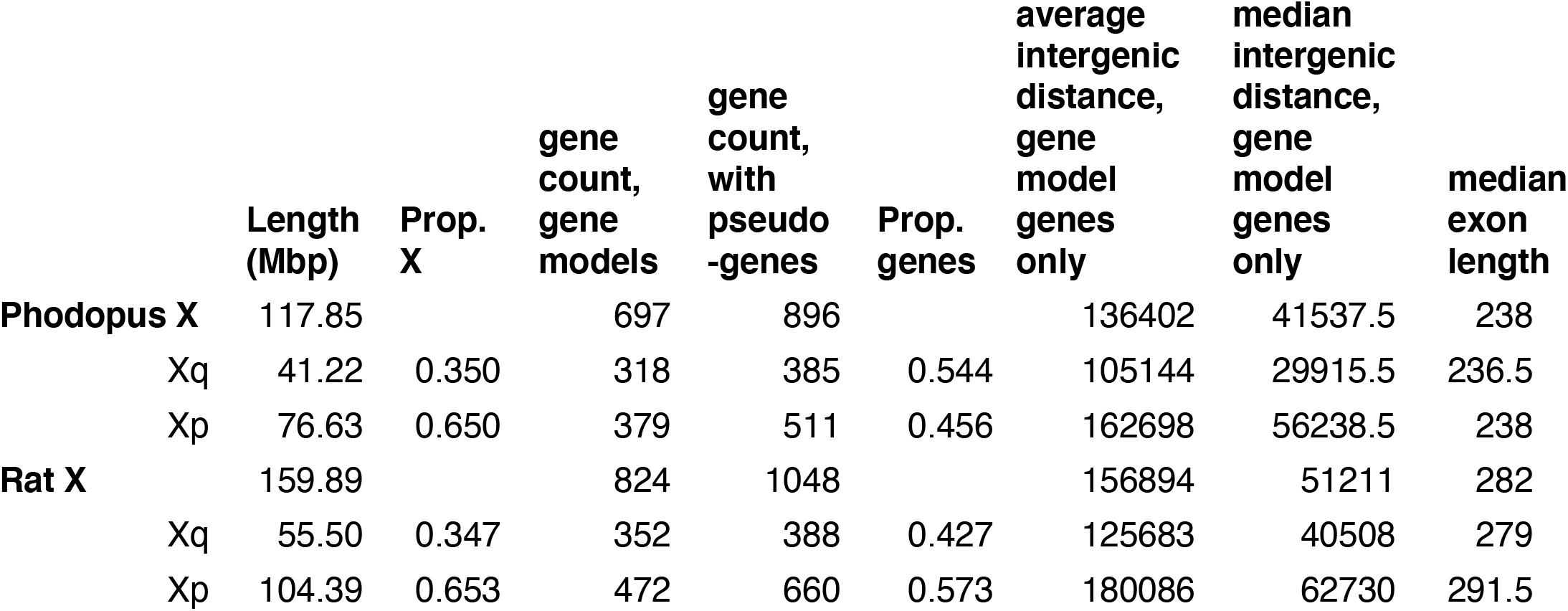
Comparison of features on the X chromosomes of dwarf hamster and rat

**Table S4.**
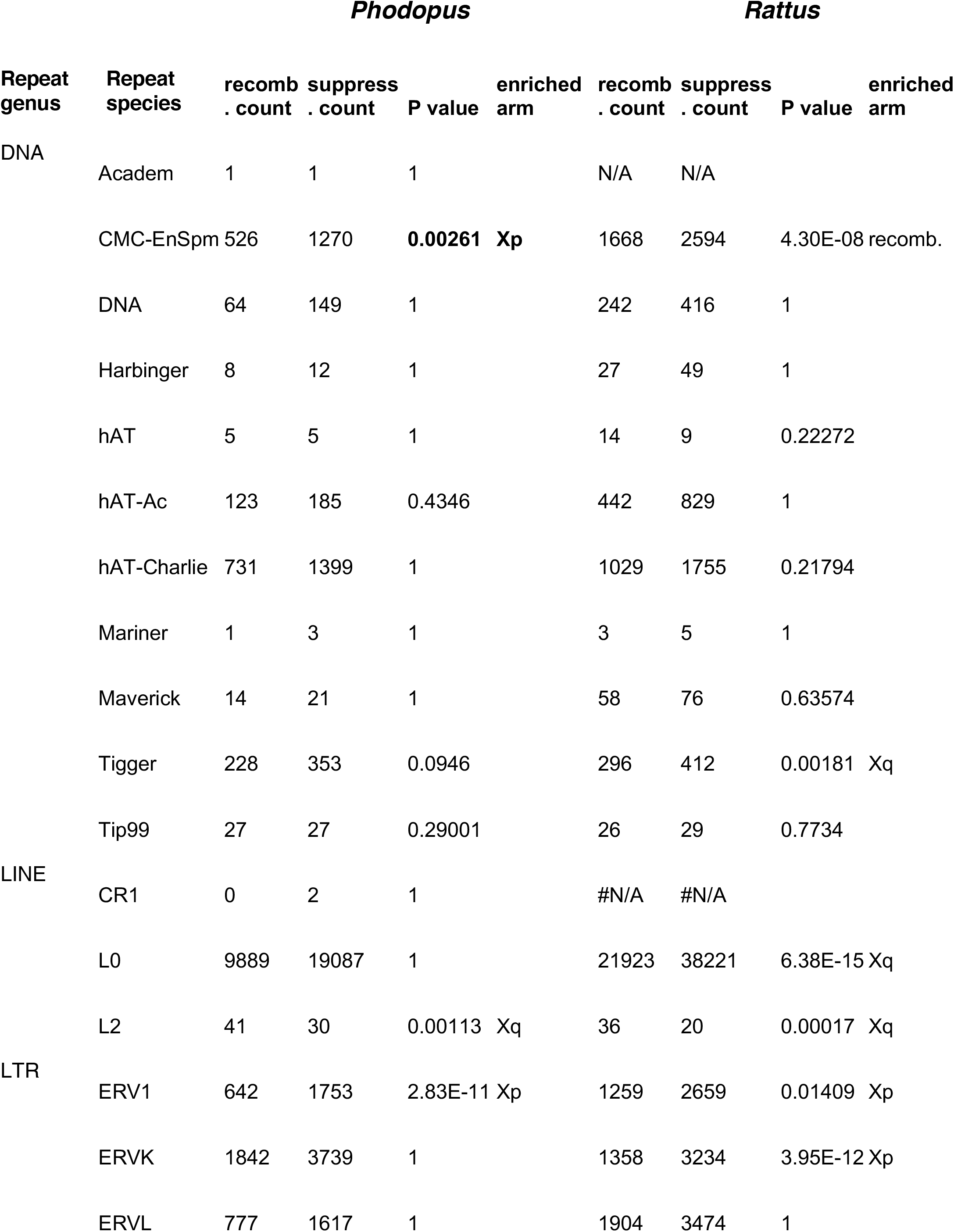
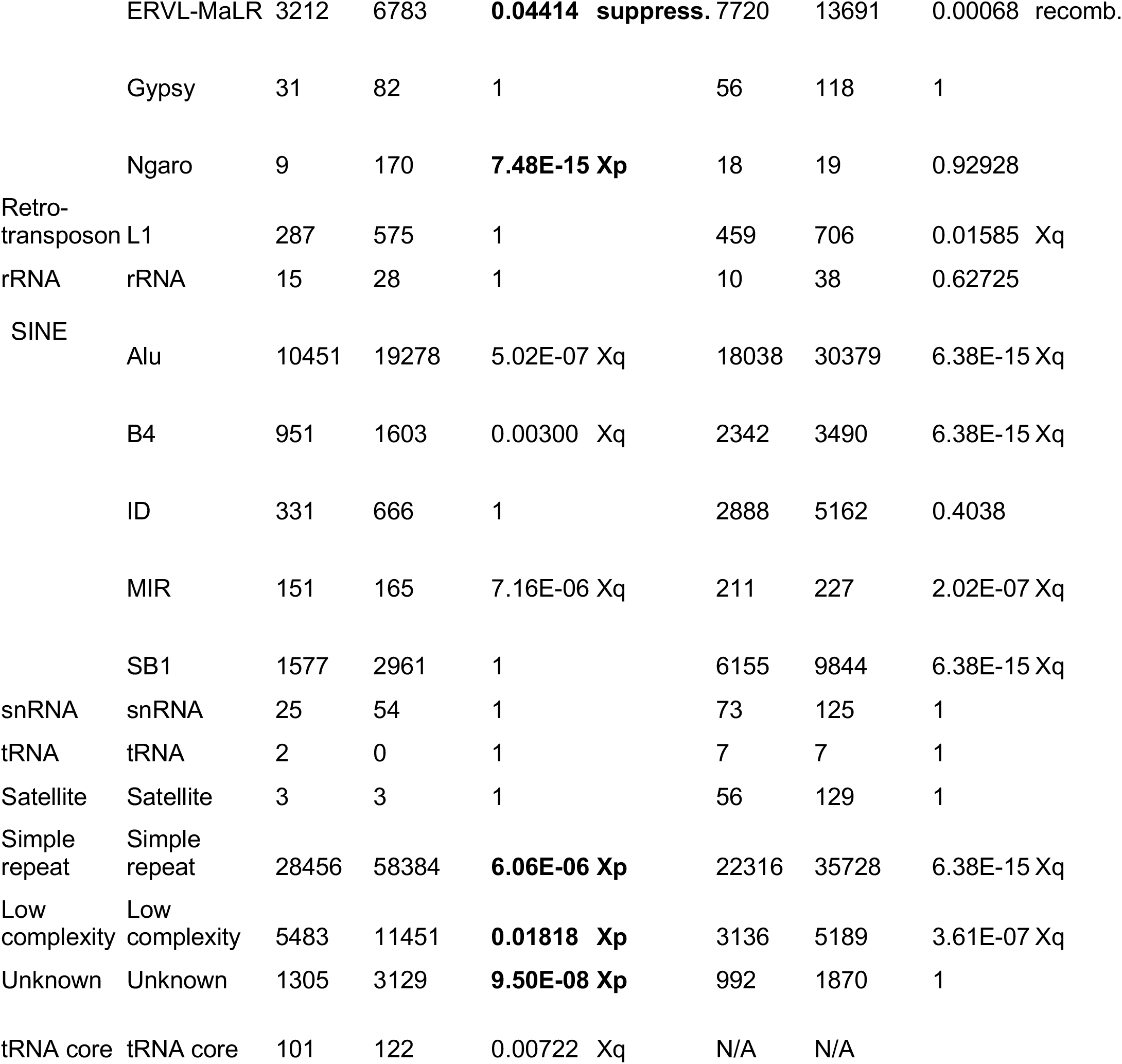
Transposable element enrichment on X chromosome arms

**Table S5:**
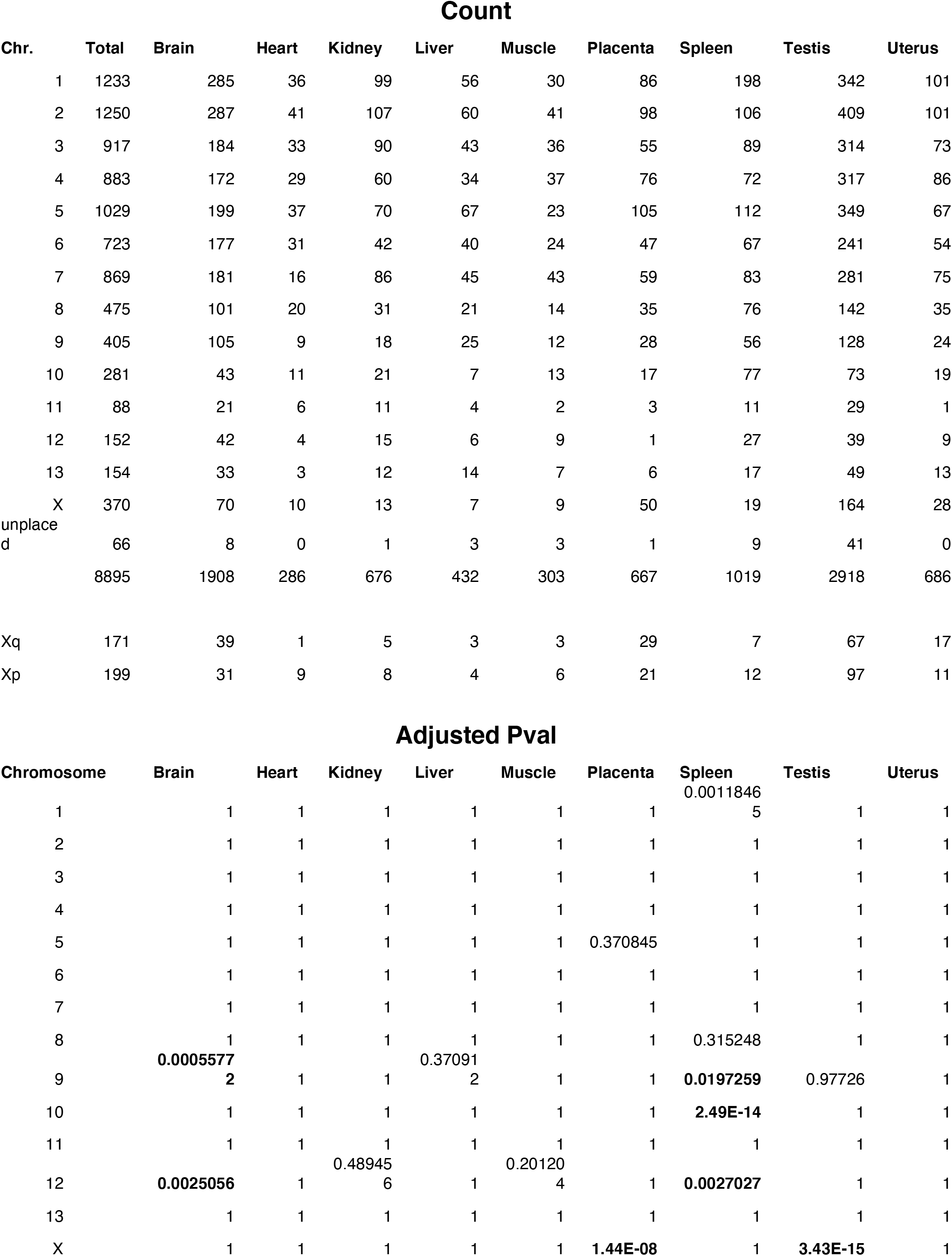
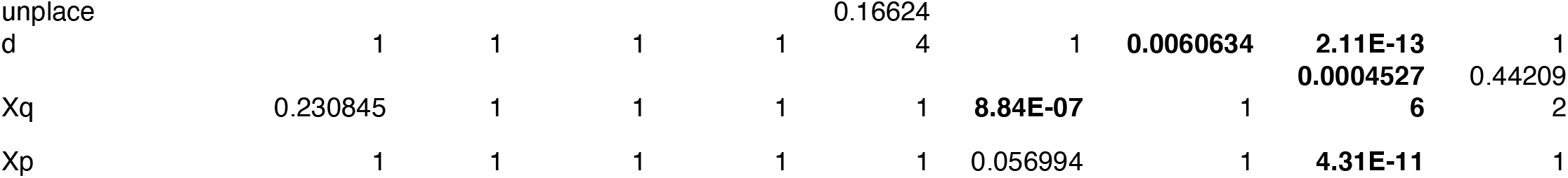
Counts of tissue-enriched genes by chromosome, and BH corrected p-values

**Supplementary Figure 1.**
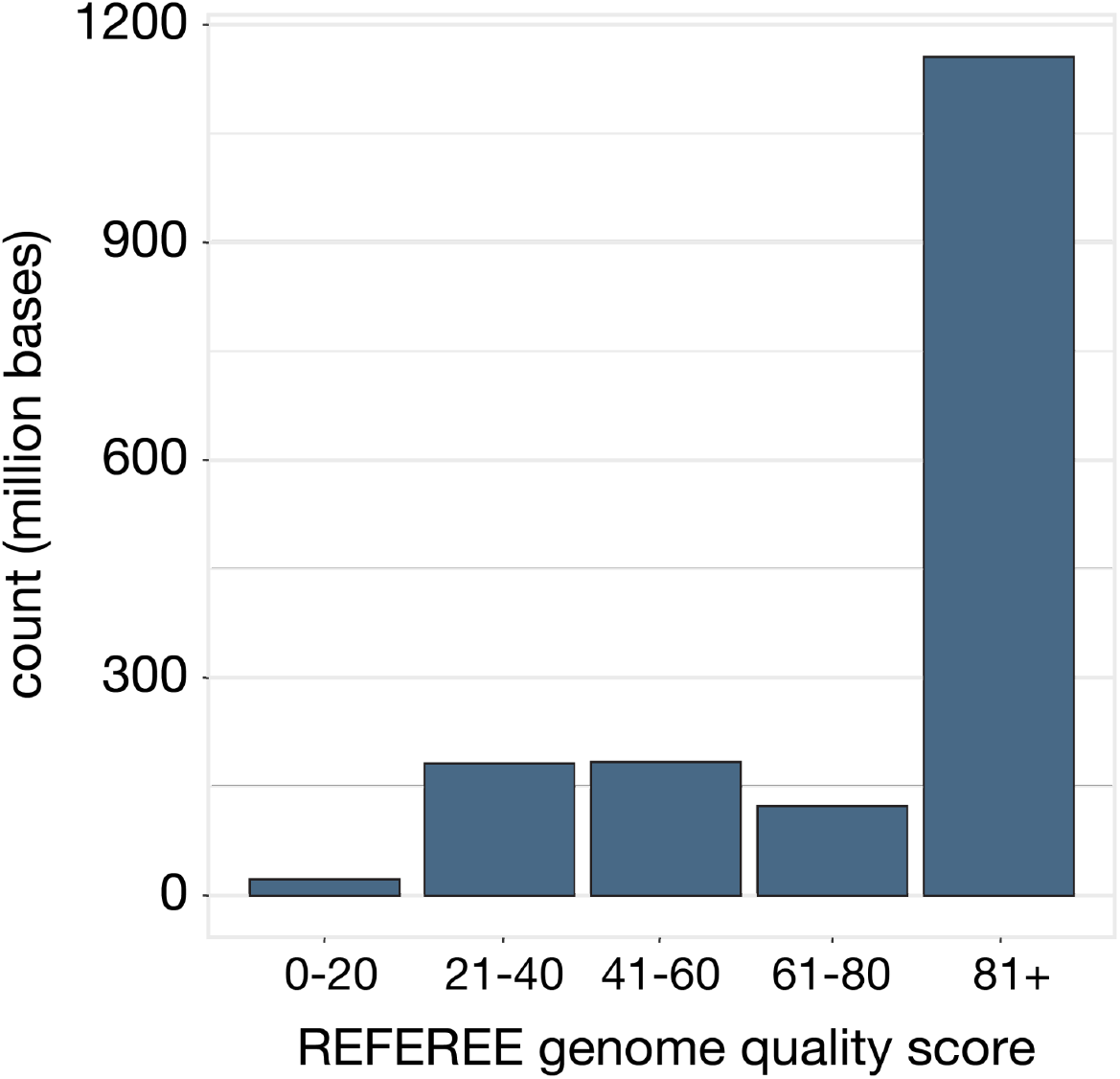
Distribution of site-based quality scores from the largest scaffold per chromosome from Referee.

**Figure S2.**
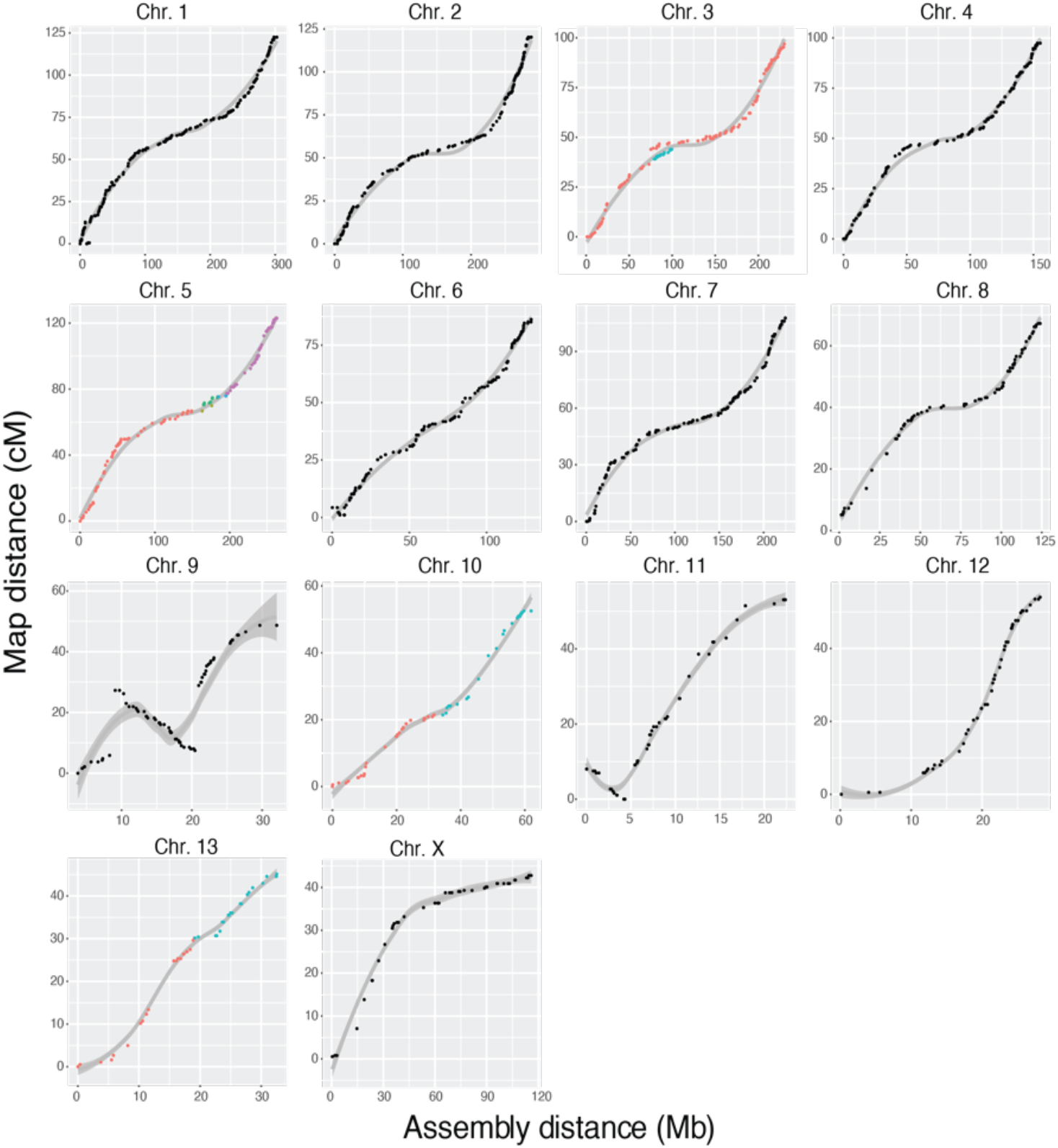
Marker positions by genetic map (cM) and assembly positions (Mb) for all chromosomes. Genetic and physical marker locations show recombination across each chromosome for all anchored scaffolds over 1Mb, with the line showing a smoothed spline best fit. The slope of this line reflects recombination rate, with a steeper slope indicating a higher recombination rate. Metacentric chromosomes 1-8, 10, and 13 show an expected reduction in recombination rate near the centromere, whereas chromosome 12 shows an initial reduction in recombination that likely reflect acrocentric centromeres. Despite being metacentric, the X chromosome never recovers recombination on the Xp arm. Regions of chromosome 9 and chromosome 13 with negative slopes likely reflect assembly errors. Colored points on chromosomes 3, 5, 10, and 13 indicate scaffolds, all other chromosomes consist of one major scaffold.

**Supplementary Figure 3.**
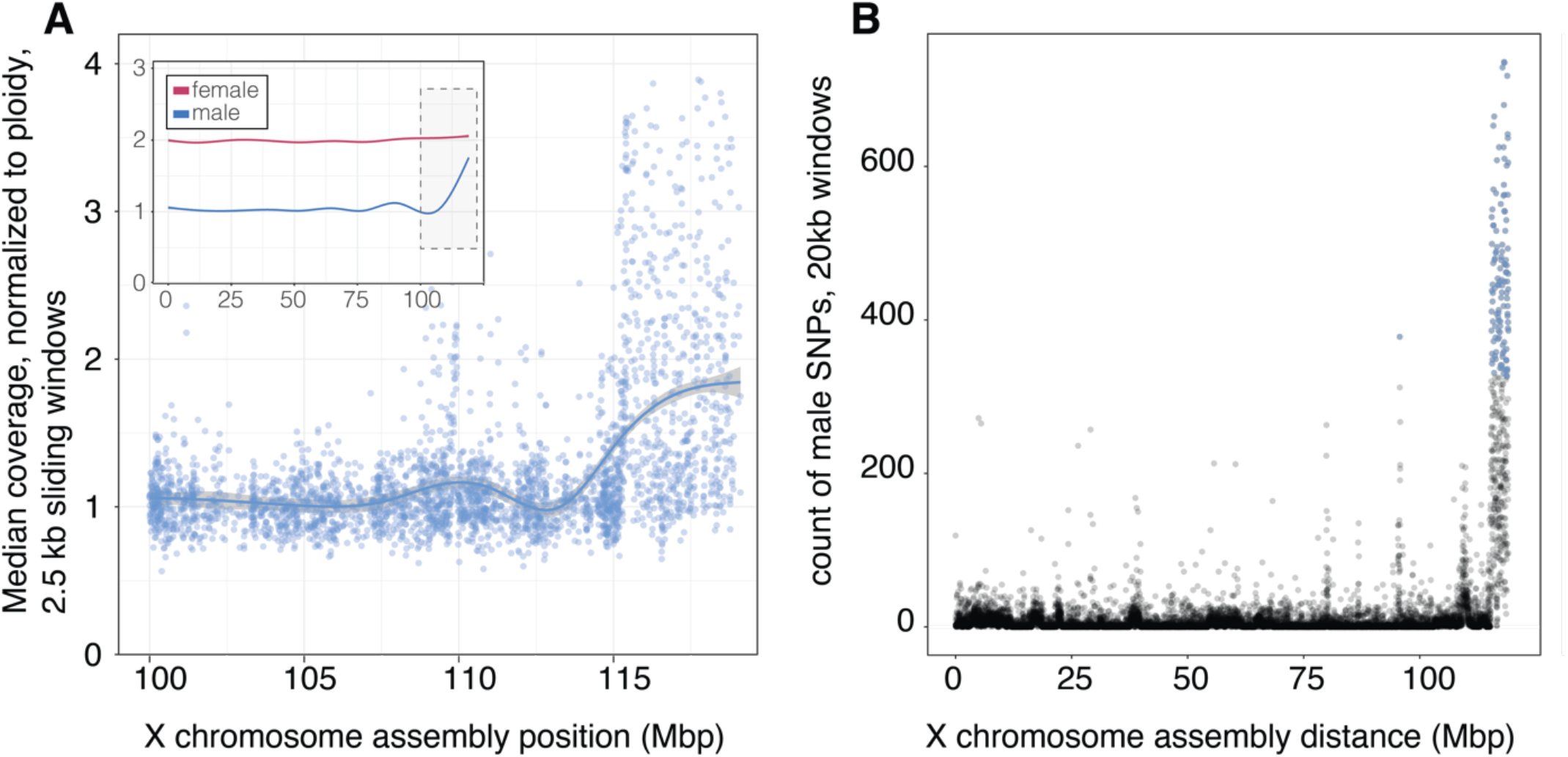
Identification of the pseudoautosomal region of the X chromosome. (A) Inset, median coverage normalized to ploidy along the entire *P. sungorus* X chromosome for males (blue) and females (female). The main panel shows the distal X chromosome, where the pseudoautosomal region is indicated by an increase in coverage where reads from the Y map to homologous sequence on the X. Mean coverage calculated in 2.5 kbp sliding windows. (B) The distal end of the chromosome also shows an increase in the number of SNPs called from the male sequence, suggesting that these reads come from divergent Y sequence. Count of SNPs in 20 kbp windows. For (B), blue points indicate windows in the top and bottom 1% of the distribution of values.

**Supplemental Figure 4.**
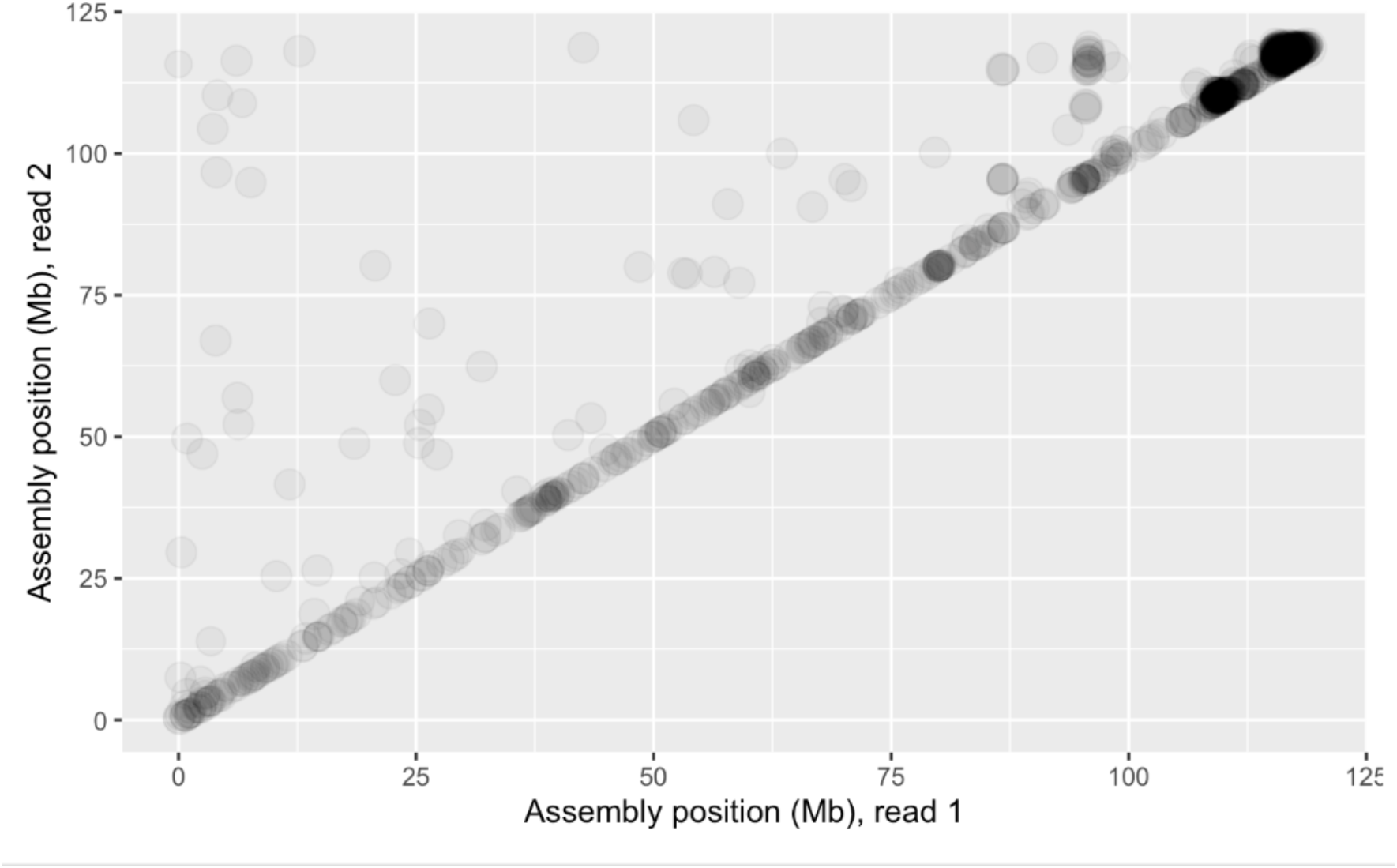
Short-read support an inversion between *P. sungorus* and *P. campbelli* on the X chromosome. Differences in the mapping location of read 1 (x-axis) and read 2 (y-axis) in a read-pair indicate an inversion in *P. campbelli*, using short-reads from *P. sungorus* as a baseline. Darker colors indicate more support for an inversion in this location.

**Supplemental Figure 5.**
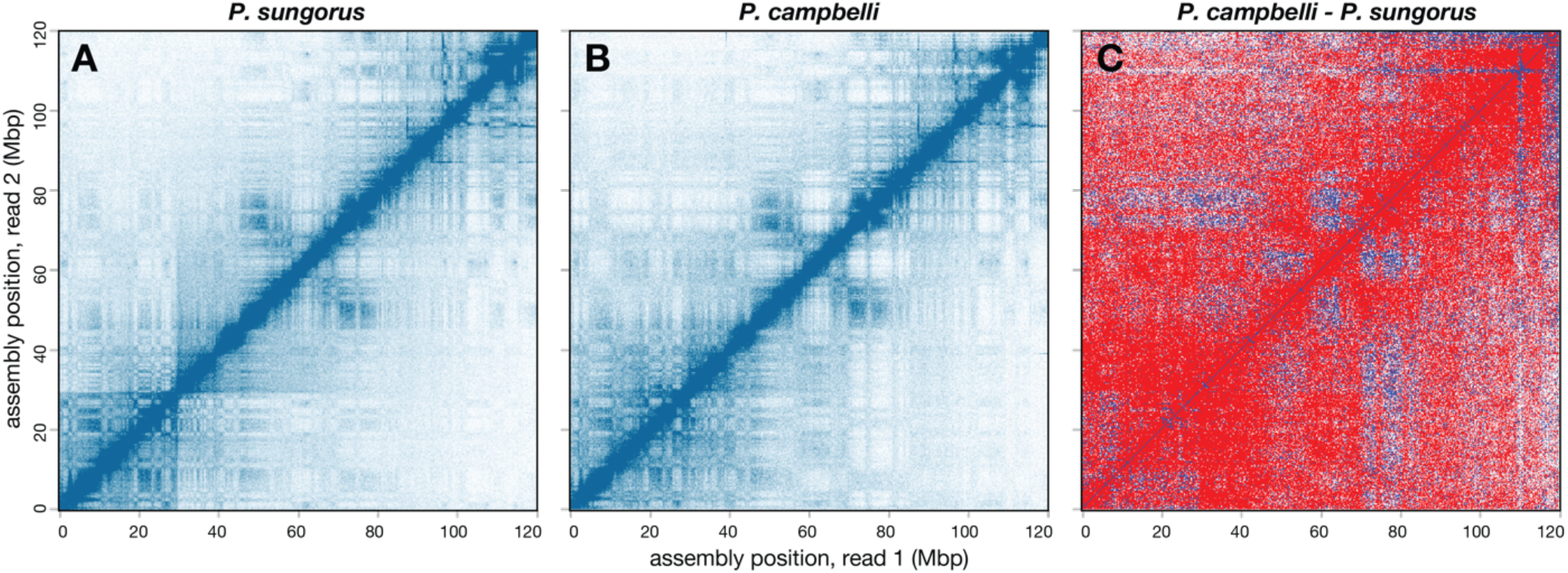
Comparative chromatin configuration of the X chromosome between *P. sungorus* and *P. campbelli*. HiC chromatin interactions, show short- and long-range interactions between points on the X chromosome, 250 Kbp resolution, square root coverage normalization for (A) *P. sungorus* and (B) *P. campbelli*. (C) The difference between the two, where blue indicates increased contact in the *P. sungorus* chromatin map relative to the *P. campbelli* map.

**Supplemental Figure 6.**
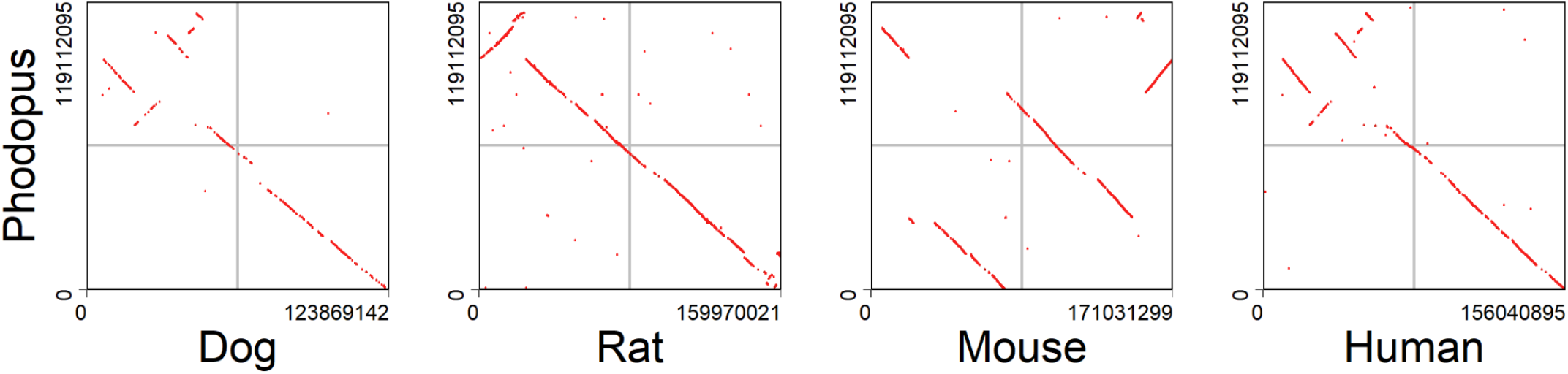
X chromosome synteny between mammalian species. Alignments of the X chromosomes of dwarf hamster to domestic dog (CanFam3.1), rat (rnor6), mouse (mm10), and human (GRCh38) show broad conservation of synteny across mammals with the exception of mouse.

**Supplemental Figure 7.**
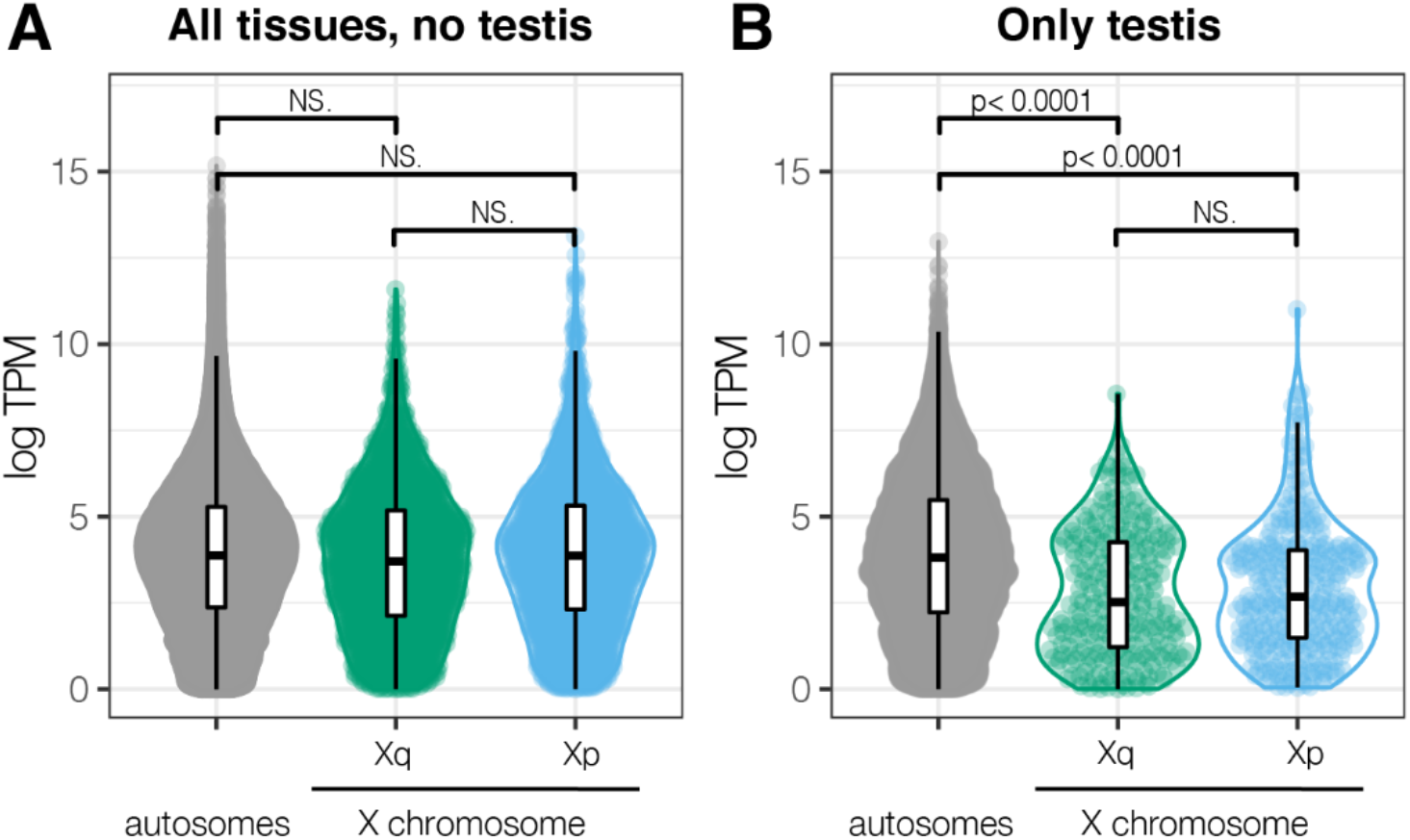
Gene expression levels on the X chromosome arms vs autosomes. Gene expression levels (log2TPM [transcripts per million] for (A) all tissues expect testis and (B) testis (significance, pairwise Wilcoxon).

**Supplemental Figure 8.**
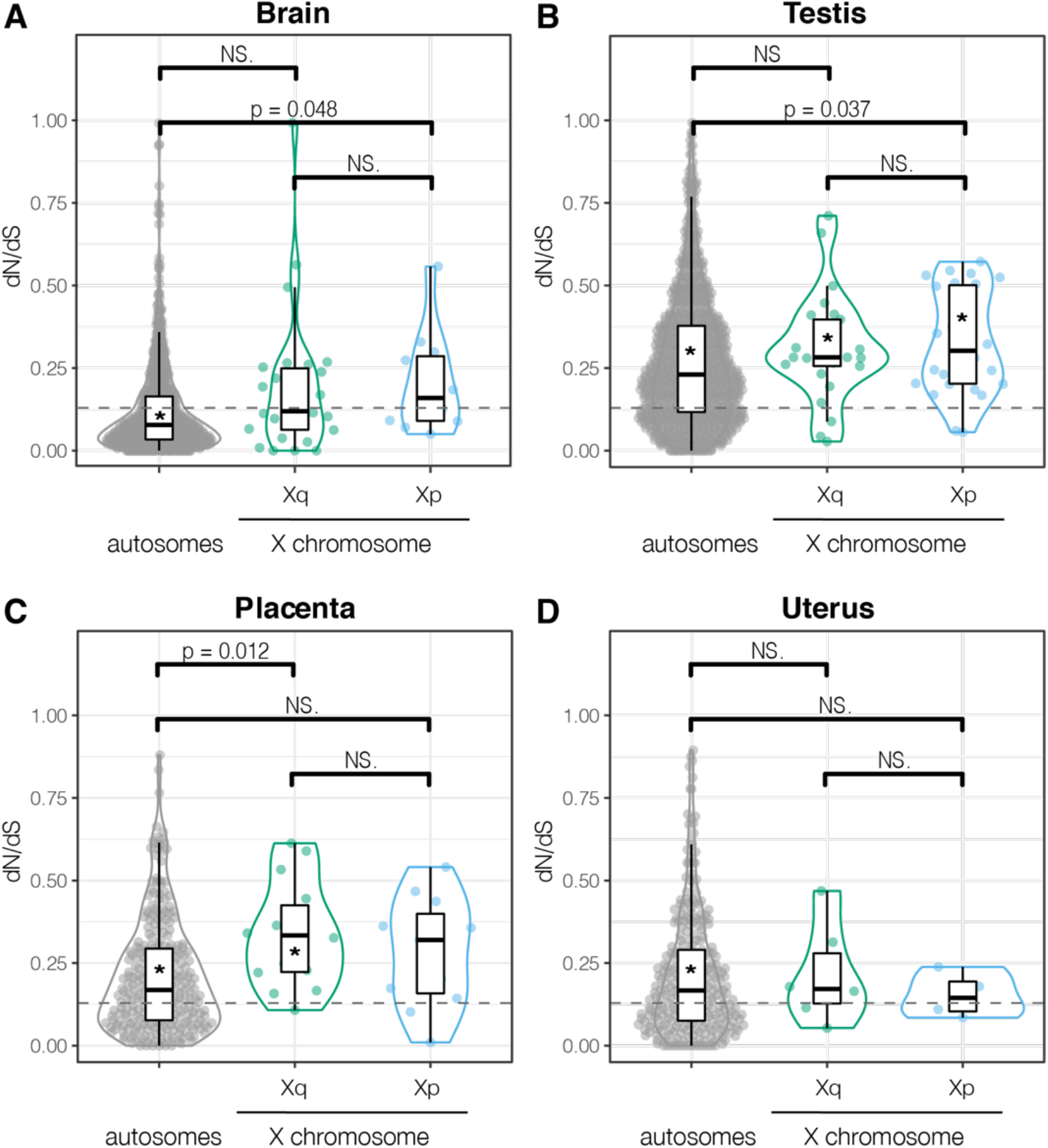
Evolutionary rates in genes with high reproductive tissue specificity. dN/dS for genes with Tau greater than 0.8, for four tissues with sufficient numbers of genes on both X chromosome arms. Significance between autosomes and X arms indicated with bars and corrected p-values (pairwise Wilcoxon). Dashed line indicates genome-wide dN/dS for all genes, all tissues; significant deviation from the genome-wide median dN/dS values is indicated with an asterisk (*) in the box plot (pairwise Wilcoxon).

**Supplemental Figure 9:**
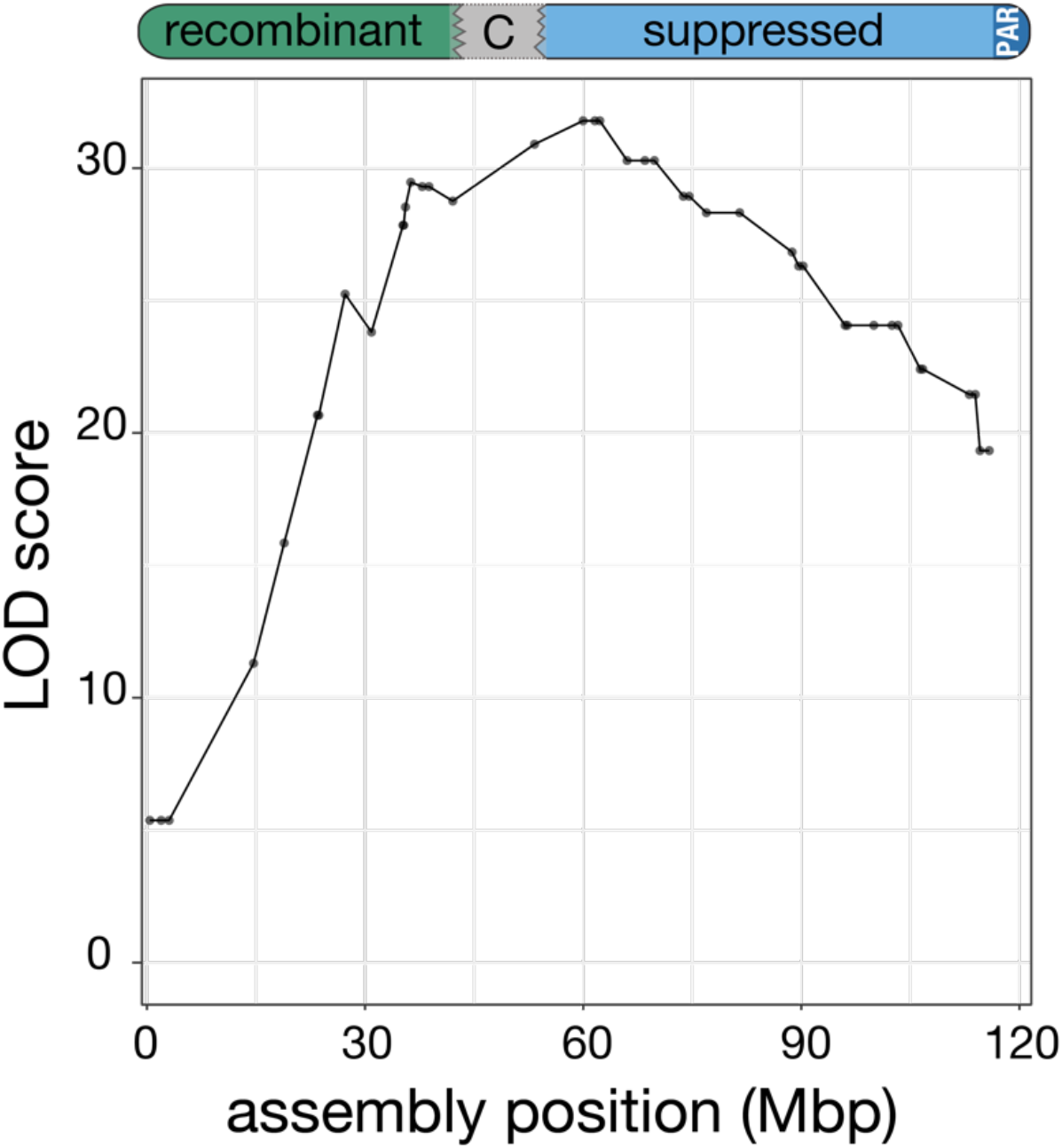
Placental hybrid incompatibility QTL position on the X chromosome genome build. Significance of association (LOD score) for X-linked hybrid incompatibility QTL shown according to marker assembly location (in Mbp), rather than genetic map (data from Brekke *et al* 2021)

**Supplemental Figure 10:**
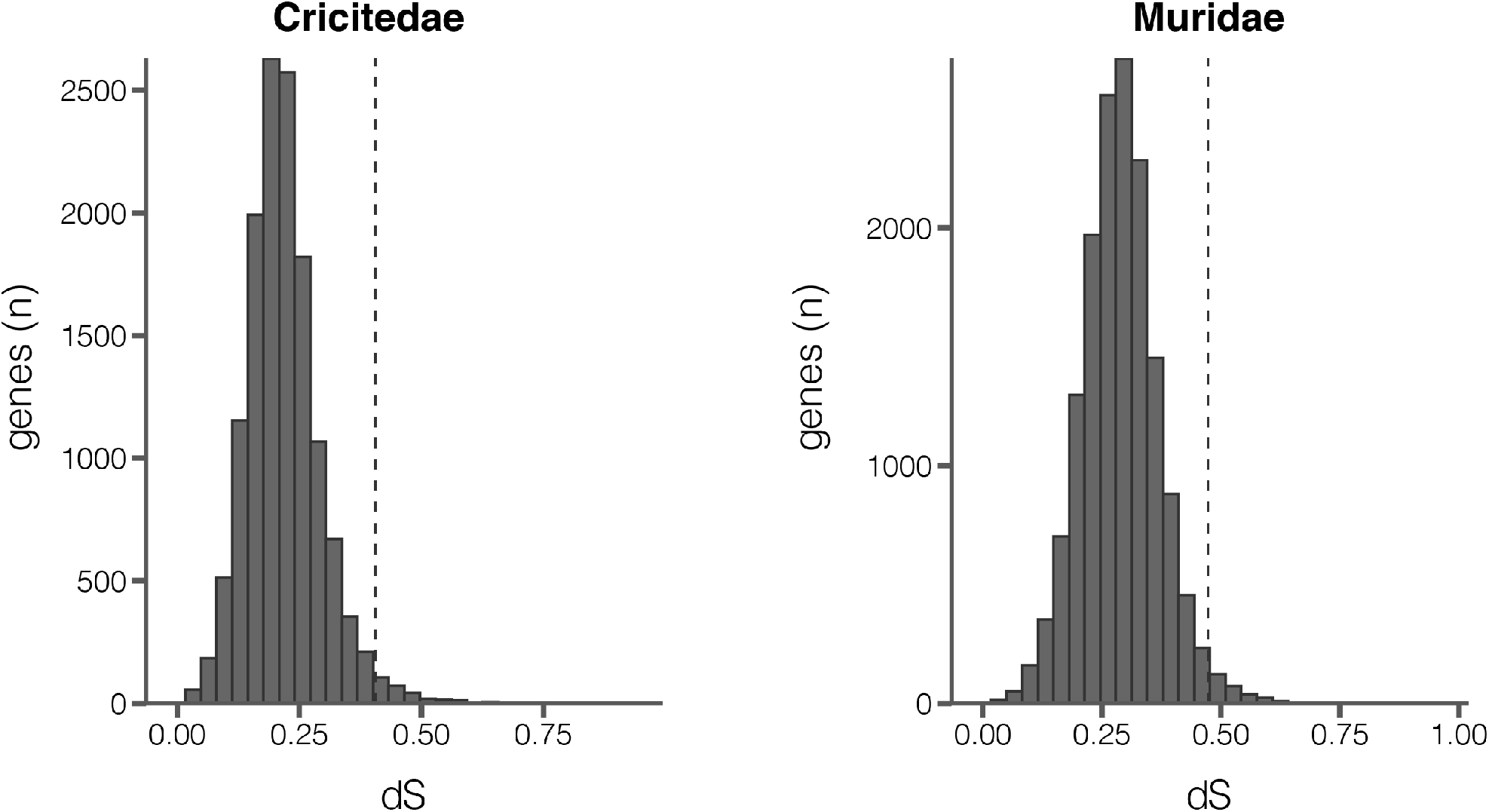
Distributions of dS among single copy orthogroups identified in four species in Cricitedae and four species in Muridae. The vertical dashed line represents the 98^th^ percentile, above which genes were removed from subsequent analysis as a control for possible alignment error.

